# Single-Cell Manifold Preserving Feature Selection (SCMER)

**DOI:** 10.1101/2020.12.01.407262

**Authors:** Shaoheng Liang, Vakul Mohanty, Jinzhuang Dou, Qi Miao, Yuefan Huang, Muharrem Müftüoğlu, Li Ding, Weiyi Peng, Ken Chen

## Abstract

A key challenge in studying organisms and diseases is to detect rare molecular programs and rare cell populations (RCPs) that drive development, differentiation, and transformation. Molecular features such as genes and proteins defining RCPs are often unknown and difficult to detect from unenriched single-cell data, using conventional dimensionality reduction and clustering-based approaches.

Here, we propose a novel unsupervised approach, named SCMER, which performs UMAP style dimensionality reduction via selecting a compact set of molecular features with definitive meanings.

We applied SCMER in the context of hematopoiesis, lymphogenesis, tumorigenesis, and drug resistance and response. We found that SCMER can identify non-redundant features that sensitively delineate both common cell lineages and rare cellular states ignored by current approaches.

SCMER can be widely used for discovering novel molecular features in a high dimensional dataset, designing targeted, cost-effective assays for clinical applications, and facilitating multi-modality integration.

## 2 Introduction

A tissue in a living organism often consists of millions to billions of cells. While the terminally differentiated cells with relatively distinct molecular profiles can be readily distinguished via single-cell RNA sequencing (scRNA-seq) at current sampling depth, many cells involved in development, differentiation, and transformation remain difficult to detect^1,2^. For example, a fraction of tumor cells in renal cell carcinomas can go through sarcomatoid transformation driven by epithelial to mesenchymal transformation (EMT)^3,4^; tumor cells in pancreatic ductal adenocarcinomas can transiently express stemness features (e.g., SOX2) at its invasion fronts^5–7^. These cells can be relatively rare in the sampled populations, transiently expressing certain molecular features and thereby may not form distinct clusters in high dimensional feature spaces^8,9^.

To detect characteristic features (e.g., genes, proteins) in a single-cell dataset, studies^8,10–13^ often employ unsupervised clustering followed by one-cluster-vs-all differential expression (DE) analysis. These approaches can detect major cell types governed by lineage features that dominate data variance, but are insensitive to rare but unique features that have relatively small variance and manifest as level gradients within cell-type clusters (a.k.a. cell states)^14^. They are also clumsy at detecting features affecting multiple clusters, e.g., transcription factors (TFs) regulating multiple cell types^15^, as that involves comparison of an exponentially growing number of cluster combinations. To detect features associated with continuous developmental processes, many studies perform trajectory inference^16^ followed by regression analysis to identify correlated features (e.g., Monocle^17^). The selection of features depends on trajectories, which could be challenging to infer accurately for complex processes. A detailed comparison was performed by RankCorr^12^ across various methods such as statistical tests, logistic regression, MAST^10^, scVI^11^, and COMET^13^.

Most existing approaches regard features as independent variables without exploring their interactions^18^. As a result, they tend to identify redundant features (e.g. CD3D, CD3E and CD3G for T cells) that are highly correlated with the inferred clusters or trajectories, but ignore novel, uncorrelated features. Some recent work such as scHOT^18^ and SCMarker^19^ started to exploit correlational patterns among co- or anti-expressing genes. However, they do not model complex interactions of more than two genes. SCMarker cannot characterize continuous cell states, and scHOT relies on the accuracy of trajectory inference.

To enhance sensitivity in detecting rare features and RCPs, many studies^20,21^ had to slice and dice data spaces in empirical, multifaceted ways^8^ or perform iterative gating^22^ and re-clustering at variable resolutions, which may lead to biased, irreproducible results.

Increasing the number and variety of molecular features^23,24^ and improving the fidelity of the measurements can help discover RCPs. However, they unavoidably increase the already high cost of experiments. To make assays cost-effective towards clinical applications, it is important to select a compact actionable set of molecular features that unbiasedly represent molecular diversity in high dimensional data. This ability is important for designing and manufacturing customized assays, e.g., 10x targeted gene expression, MissionBio Tapestri^25^ and NanoString GeoMx^26^, which perform multi-omics measurements of hundreds of selected DNA, RNA, and proteins.

To address these fundamental challenges, we developed SCMER (Single-Cell Manifold Preserving Feature Selection), which selects an optimal set of features such as genes or proteins from a single-cell dataset. Similar to t-Distributed Stochastic Neighbor Embedding (t-SNE)^27^ and Manifold Approximation and Projection (UMAP)^28^, we hypothesize that a manifold defined by pairwise cell similarity scores sufficiently represents the complexity of the data, encoding both global relationship between cell groups and local relationship within cell groups^29^. By preserving such a manifold while performing feature selection, the most salient features that unbiasedly represent the original molecular diversity will be selected.

SCMER does not require clusters or trajectories, and thereby circumvents the associated biases. It is sensitive to detect diverse features that delineate common and rare cell types, continuously changing cell states, and multicellular programs^15^ shared by multiple cell types. It reduces high dimensionality into a compact set of actionable features with definitive biological meanings. This distinguishes SCMER from PCA, t-SNE, UMAP, etc., which result in axes (meta-genes) with complex meanings. SCMER is efficiently implemented in Python using PyTorch^30^,and supports multicore and GPU acceleration. A one-liner interface is provided for easy use. The open-source implementation is available at https://github.com/KChen-lab/SCMER.

## 3 Results

### 3.1 The SCMER Approach AND ITS Unique Strength

In a nutshell, SCMER (**Fig. 1a, Methods**) examines a data matrix **x** of *n* cells and ***D*** features and calculates a pairwise cell similarity matrix **P**(as defined in UMAP^28^) representing the manifold in **X**. It defines a weight vector **w**, which transforms **X** to **Y= Xw**. It then calculates another pairwise cell similarity matrix **Q** from **Y** and quantifies the level of agreement between **P** and **Q** using Kullback-Leibler (KL) divergence as defined in t-SNE^27^. Finally, it uses elastic net to find a sparse and robust solution of **w** that minimizes the KL-divergence with Orthant-Wise Limited Memory Quasi-Newton (OWL-QN) algorithm^31^. Features with nonzero weights in **w** are deemed chosen. **Q** can also be calculated from a different source (e.g., a different technology) instead of **X**, which enables a “supervised” and multi-omics mode of SCMER.

**Fig 1:**
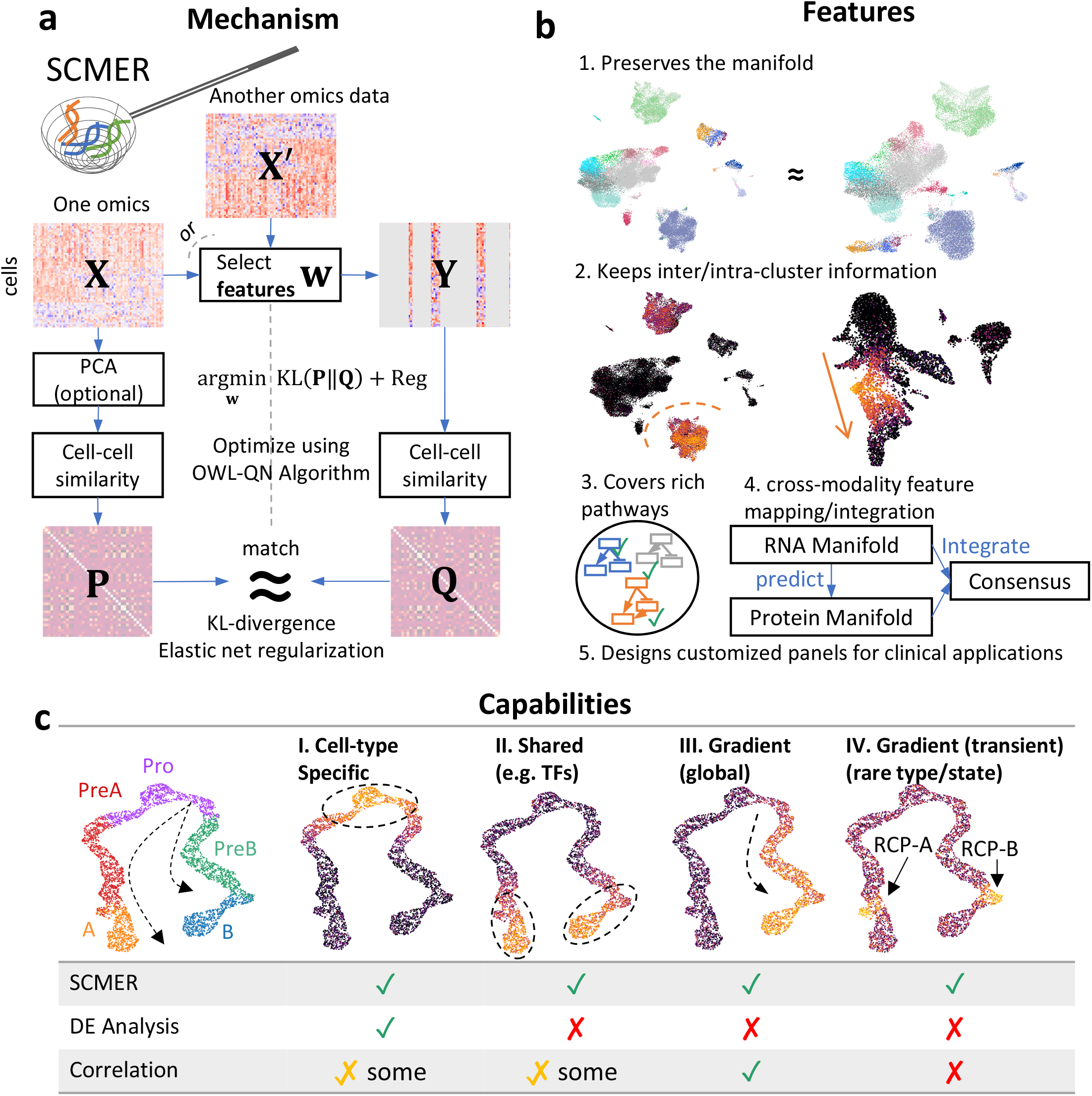
The SCMER approach and its unique strength. **(a)** Workflow of SCMER. SCMER selects the features that perserve the manifold from a single-cell omics dataset ***X***. Features can be selected from either **X** or another co-assayed omics **X**′. Vector ***w*** indicates the selection. **Y** is the dataset after feature selection. Pand Q are cell-cell similarity matrices for **X** and Y, respectively. **(b)** Applications of SCMER. SCMER selects features that preserve the manifold and retain inter- and intra-cluster diversity, and thus can be applied to discover rich molecular pathways, integrate modalities, and design customized DNA/RNA/antibody panels of restricted sizes. **(c)** Capabilities of SCMER compared with mainstream label/cluster-based differential expression (DE) analysis methods and correlation-based methods. The hypothetical branching trajectories contains common progenitors (Pro), precursors (PreA, PreB), and mature cells (A, B).

A manifold encodes both clusters and continuums of cells. While clusters usually reflect distinct cell types, continuums reflect similar cell types and trajectory of transitioning/differentiating cell states^32^. SCMER selects optimal features that preserve the manifold and retain inter- and intra-cluster diversity (**Fig. 1b**). It can be applied to discover rich molecular pathways, identify prognostic genes, and design customized DNA/RNA/antibody panels of restricted sizes towards clinical applications.

To elucidate the novel cell populations and features that SCMER uniquely identifies, we generated a synthetic dataset containing a branching trajectory of 4,000 single cells from five major cell types, namely progenitor (Pro), precursor of A and B (PreA and PreB), and mature A and B (A and B) (**Fig. 1c**). Four kinds of features are simulated, those (I) specific to one cell type/cluster, (II) shared by more than one cell type^15^, (III) gradually changing over cell states, and (IV) transiently activated (also known as checkpoints^33^). We created a total of 180 features including 20 cell-type specific features for each cell type (100 in total), 10 gradually changing features for each branch (20 in total), 5 shared features for precursors and mature cells (10 in total), 5 transiently expressing features, and 45 random noise features. Each cell was given a ground-truth pseudo-time in the trajectory, which parameterizes the expected level of a feature. In addition to major cell type labeling, the cells transitioning from precursor to mature are identified as “RCP-A” and “RCP-B”, which overexpress type-IV features. Dispersion was added based on a negative-binomial distribution. In as few as 45 selected features, SCMER recalled all the four types of features. In contrast, the top 45 features determined by a DE analysis are all from type I -- none belonged to type II, III, or IV, while a pseudo-time-based correlation analysis missed type-IV features. As a result, SCMER significantly increased the precision and recall of detecting RCPs, while being comparable to other methods on major cell types (using a *k*-NN classifier for one cell type at a time; **Table 1** and **Supplementary Note 1**).

**Table 1.** Precision/recall of detecting RCPs on the synthetic dataset

| Cell types | RCPs |  | Major cell types |  |  |  |  |
| --- | --- | --- | --- | --- | --- | --- | --- |
|  | RCP-A | RCP-B | Pro | PreA | PreB | A | B |
| Abundance | 2.55% | 2.68% | 21.23% | 22.43% | 19.83% | 16.73% | 15.30% |
| SCMER | 0.82 / 0.68 | 0.87 / 0.67 | 0.97 / 0.96 | 0.95 / 0.96 | 0.94 / 0.94 | 0.95 / 0.96 | 0.94 / 0.93 |
| DEG | 0.61 / 0.34 | 0.73 / 0.40 | 0.94 / 0.95 | 0.94 / 0.93 | 0.95 / 0.94 | 0.94 / 0.95 | 0.95 / 0.96 |
| Correlation | 0.48 / 0.36 | 0.43 / 0.28 | 0.91 / 0.96 | 0.76 / 0.67 | 0.76 / 0.67 | 0.88 / 0.95 | 0.88 / 0.92 |

To comprehensively assess SCMER, we ran it on eight datasets^34–41^ (including **Supplementary Results**) that involve a variety of biological and technological challenges, such as unresolved borderline cells that blur clustering, continuously changing cell states, multicellular and transient cellular programs. For comparison, we used supervised DE analysis and widely-used unsupervised feature selection methods, including highly expressed genes (HXG), highly variable genes (HVG), SCMarker^19^, Monocle^17^, and RankCorr^12^. SCMER robustly demonstrated the best performance on all the experiments.

### 3.2 SCMER CHARACTERIZATION OF CELL TYPE AND INTRATUMORAL HETEROGENEITY IN CANCER

Single-cell datasets derived from cancer samples are often highly complex, containing heterogeneous cell types and states in not only tumor cells but also stromal and immune cells. Supervised analysis of cancer data is challenging as cancer cells are highly plastic^42^ and can express novel unknown features, which can heavily confound clustering and trajectory-based analysis. We applied SCMER on a single-cell RNA-seq melanoma dataset containing 4,645 cells from 19 human melanoma samples^34^. Most cells were annotated as malignant cells, B cells, T cells, macrophages, natural killer (NK) cells, endothelial cells, or cancer associated fibroblasts (CAFs) by the authors based on clustering and DE analysis. However, there were unresolved borderline cells presenting between labeled clusters, which resemble more than one cell types and could be either doublets or RCPs (**Fig. 2a**). By selecting only 75 genes (**Table 2**), SCMER clearly preserved the manifold: the resulting UMAP embedding is very similar to the original and the relations among most cell types including the unresolved cells are largely preserved (**Fig. 2b** and **Supplementary Figure 2**).

**Table 2.** Melanoma features selected by SCMER

| Category | Gene set | All | Highly variable | SCMER | Genes |
| --- | --- | --- | --- | --- | --- |
| Resistance / intratumor heterogeneity | AXL | 100 | 52 | 2 | *HAPLN3, CD52* |
|  | MITF | 100 | 28 | 3 | *TOB1, TYR, PMEL* |
| Cell types | Melanoma | 47 | 19 | 5 | *SERPINA3, PRAME, MIA, TYR, PMEL* |
|  | B cells | 31 | 27 | 2 | *IRF8, MS4A1* |
|  | T cells | 38 | 38 | 7 | *TCF7, TIGIT, CD8A, PRDM1, SPOCK2, NKG7, TOX* |
|  | Macrophages | 92 | 73 | 1 | *TYROBP* |
|  | Endothelial | 95 | 45 | 2 | *CDH5, HAPLN3* |
|  | Cancer associated fibroblasts (CAF) | 88 | 46 | 3 | *COL1A2, COL1A1, MFAP4* |
| Pathways | Exhaustion (consistent across samples) | 28 | 17 | 3 | CD27, TIGIT, CXCL13 |
|  | Exhaustion (variable across samples) | 272 | 159 | 7 | *HLA-DPA1, HAVCR2, CTLA4, SRGN, IRF8, PDCD1, TOX* |
|  | Cell cycle | 93 | 54 | 3 | *TYMS, AURKB, MCM5* |
| None of above | *C10orf54, A2M, RGS10, MIR4461, IFITM1, SYTL3, GZMB, DEDD, RGS1, MAGEC2, PTPRC, NME2, CCL4, CCR7, HLA-DQA1, GNLY, OAS2, TXNIP, SELPLG, PAEP, CD69, ELK2AP, GZMH, ANXA1, MTRNR2L2, B2M, PTPRCAP, HLA-DQB1, LSP1, C1orf56, TCL1A, GPR183, CXCR4, RPS4Y1, APOD, KIT, GZMA, VIM, LTB, RPS11, TNFAIP3, CD55, UXS1* |  |  |  |  |

**Fig 2:**
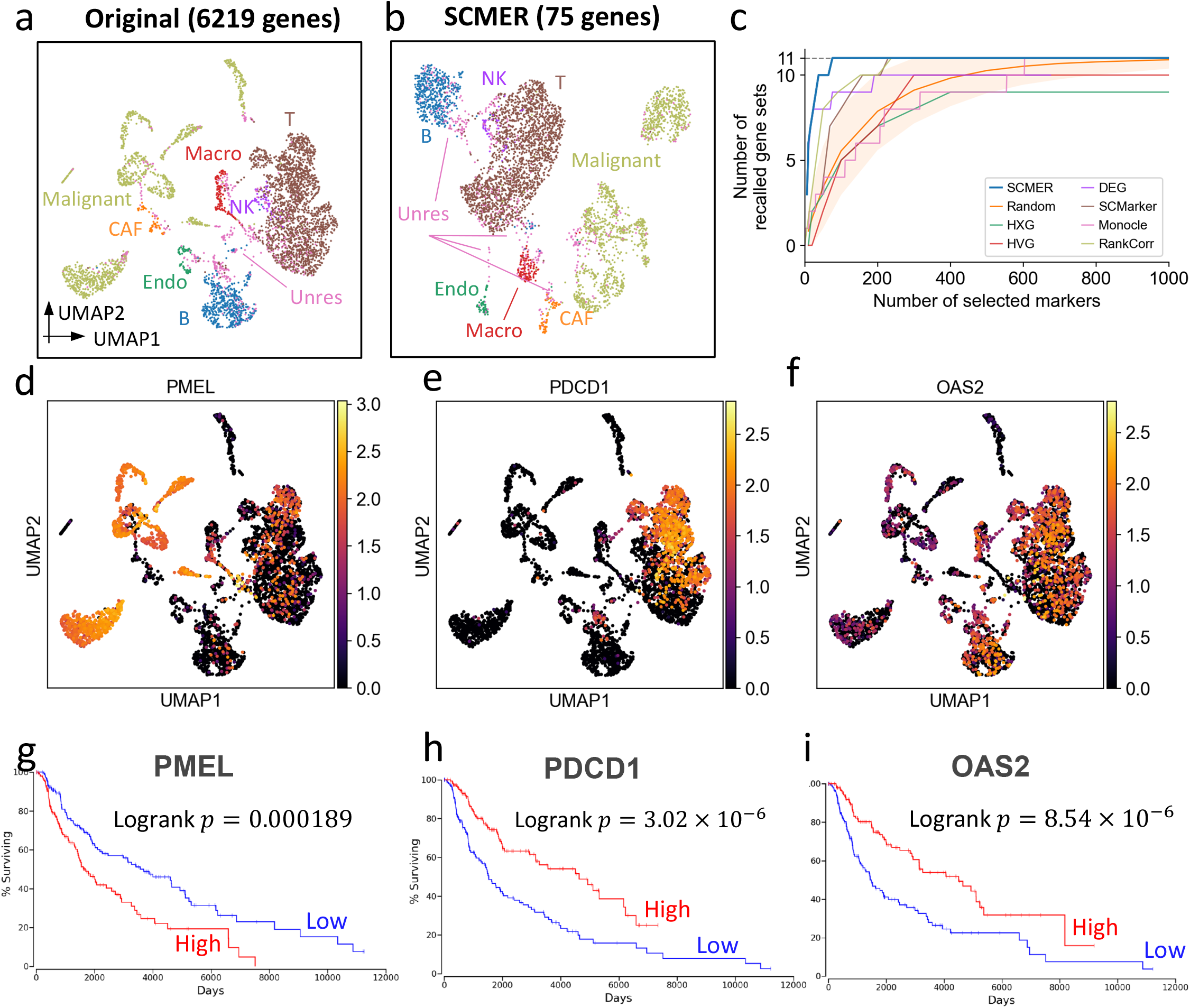
Results of the data of melanoma patients. **(a)** UMAP embedding of the dataset without feature selection. (Macro: macrophages, Endo: endothelial cells, CAF: cancer associated fibroblasts, Unres: unresolved cells; labels are in the same color of dots representing cells.) **(b)** UMAP of the dataset using SCMER selected genes. **(c)** Recall of gene sets for SCMER, scMarker, Monocle, RankCorr, highly expressed genes (HXG), highly variable genes (HVG), principal component analysis (PCA), and differentially expressed genes (DEG, supervised). X-axis is number of selected genes and Y-axis is number of covered gene sets. A gene set is recalled when at least one gene in the set is selected. Methods recalled more gene sets with fewer genes are of better performance. “Random” shows the expected number of gene sets for randomly selected markers. The area corresponds to 1.645 × standard deviation on each side. Results above the area is significantly better than random (*p<* 0.05 for one sided z-test). (**d**-**f**) Expression of genes that show intra-cluster gradients. (**g**-**i**) Overall Kaplan-Meier survival curve for selected markers in TCGA SKCM. High and low include patients in above and under 33% percentile, respectively. Each group includes *n* = 151 patients.

To understand the biological meanings of the selected genes, we compared them with the 11 gene sets described in the original publication that represent important cell types and pathways in the study. The selected genes compactly covered all the 11 gene sets (**Table 2**).

Interestingly, genes belonging to the known drug resistance AXL program and MITF program were also selected by SCMER. These genes do not preferentially express in a specific cluster (e.g. *PMEL, TOB1*, etc. in **Fig. 2d** and **Supplementary Figure 1a,b**).

SCMER also selected novel genes such as *TNFAIP3, VIM, COL1A1*, and *COL1A2*, which are involved in EMT. The original publication missed *TNFAIP3* and *VIM* and did not mention EMT. *COL1A1* and *COL1A2* were only reported as CAF specific markers. Also found were intra-cluster features for different kinds of immune cells (**Fig. 2e,f** and **Supplementary Figure 1c**) or shared by all cell types (**Supplementary Figure 1d**), which correspond to multicellular programs. *PDCD1* and *TCF7*, which are known to mark different subsets of T cells (exhausted versus memory) were detected, showing anti-correlated (Pearson’s s *r* = – 0.24) expressions within the same cluster (**Supplementary Figure 1c**). Pathway enrichment analysis^43^ on the novel genes suggest that these genes are involved in innate immune response (*A2M, CD55, IFITM1, VIM, OAS2*, etc.), adaptive immune response (*CD55, CCR7, GPR183, TNFAIP3, PTPRC*, etc.), inflammatory abnormality of the skin (*KIT, CXCR4, TCL1A, B2M*, etc.), and other immune pathways (**Supplementary Tables 1** and **2**). These genes thus may be useful in further stratifying malignant and immune cell states (e.g., activation, exhaustion, etc.). Some genes such as *PMEL, PDCD1*, and *OAS2* appeared associated with survival outcome in TCGA SKCM patients^44^ (**Fig. 1g,h,i**).

To comprehensively assess the performance of SCMER, we varied the number of selected features and recorded the number of recalled gene sets. SCMER consistently recalled more gene sets than other widely-used methods for any given number of features, demonstrating the best performance (**Fig. 2c**).

We also applied SCMER to a large-scale pan-cancer single-cell transcriptomic study consisting of 198 cell lines and patient samples from 22 cancer types^35^. SCMER showed the high sensitivity to characterize intra-cluster heterogeneity, identifying recurrent heterogeneous programs shared by a majority of cell lines and by patient tumor samples (**Supplementary Figure 3** and **Supplementary Result 1**).

**Fig 3:**
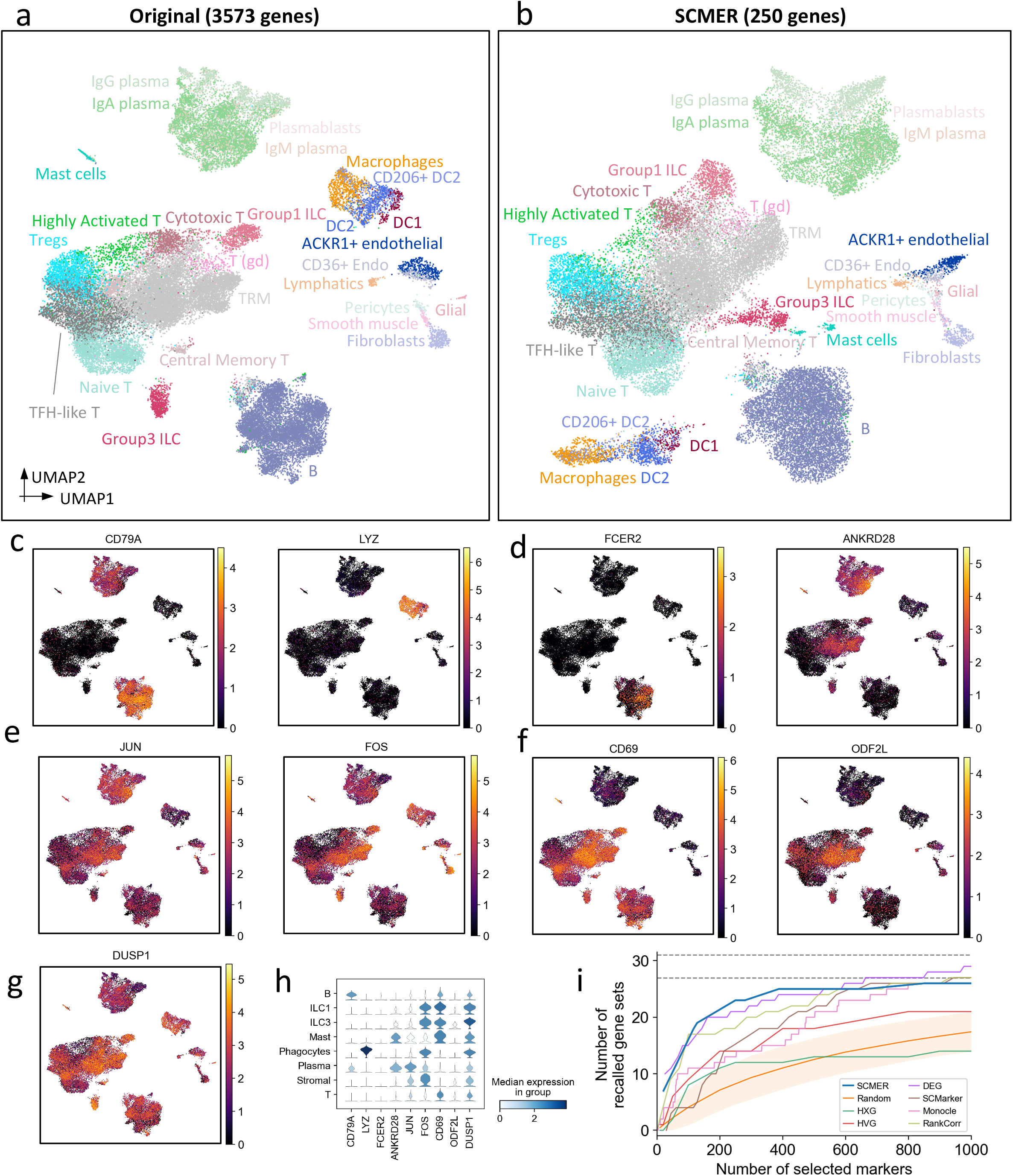
Results of the ileum lamina propria immunocytes data. **(a)** UMAP embedding of the original dataset (List of abbreviations: **Supplementary Table 14**). **(b)** UMAP embedding of the same dataset on genes selected by SCMER. (**c**-**f**) Examples of expression level of genes selected by SCMER that (c) distinguish major cell types and (**d**) subtypes, (e) are transcription factors regulating different cell types, and (f) show gradual changes among cell states. **(g)** Expression level of *DUSP1*. See **Supplementary Figure 4** for *DUSP2* and *DUSP4*. **(h)** Distribution of expressions in major cell types of genes above. **(i)** Recall of gene sets for SCMER, scMarker, Monocle, HXG, HVG, PCA, and DEG, similar to **Fig. 2c**.

### 3.3 SCMER DEFINING CELL SUBTYPES AND STATES IN ILEUM LAMINA PROPRIA IMMUNOCYTES

We further examined SCMER in a complex setting involving many cell subtypes and subtle intra-cluster structure and shared pathways. The dataset contains 39,563 gastrointestinal immune cells collected from inflamed tissues from ten Crohn’s disease patients^36^. As a risk factor to cancer, chronic inflammation involves extensive interaction among various immune cell types such as helper T cells (T_H_) and innate lymphoid cells (ILCs), which are regulated by both shared and cell-type specific TFs and cytokines and are difficult to delineate in high dimensional embeddings. The dataset appeared to include 27 cell types and subtypes/states in the original report. Four major cell types, T cells, B cells, phagocytes, and stromal cells each appeared as a cloud in the original embedding (**Fig. 3a**) but can be further dissected into subtypes. For example, T cells were dissected into eight subtypes/states through further clustering.

Circumventing clustering, SCMER selected 250 features from 3,573 highly variable genes (**Supplementary Tables 3 and 4**) with the manifold well preserved. The separability among cell types were comparable with the original embedding, and the manifold of subtypes (RCPs) in each major cell type were well preserved (**Fig. 3b**).

**Table 3.**
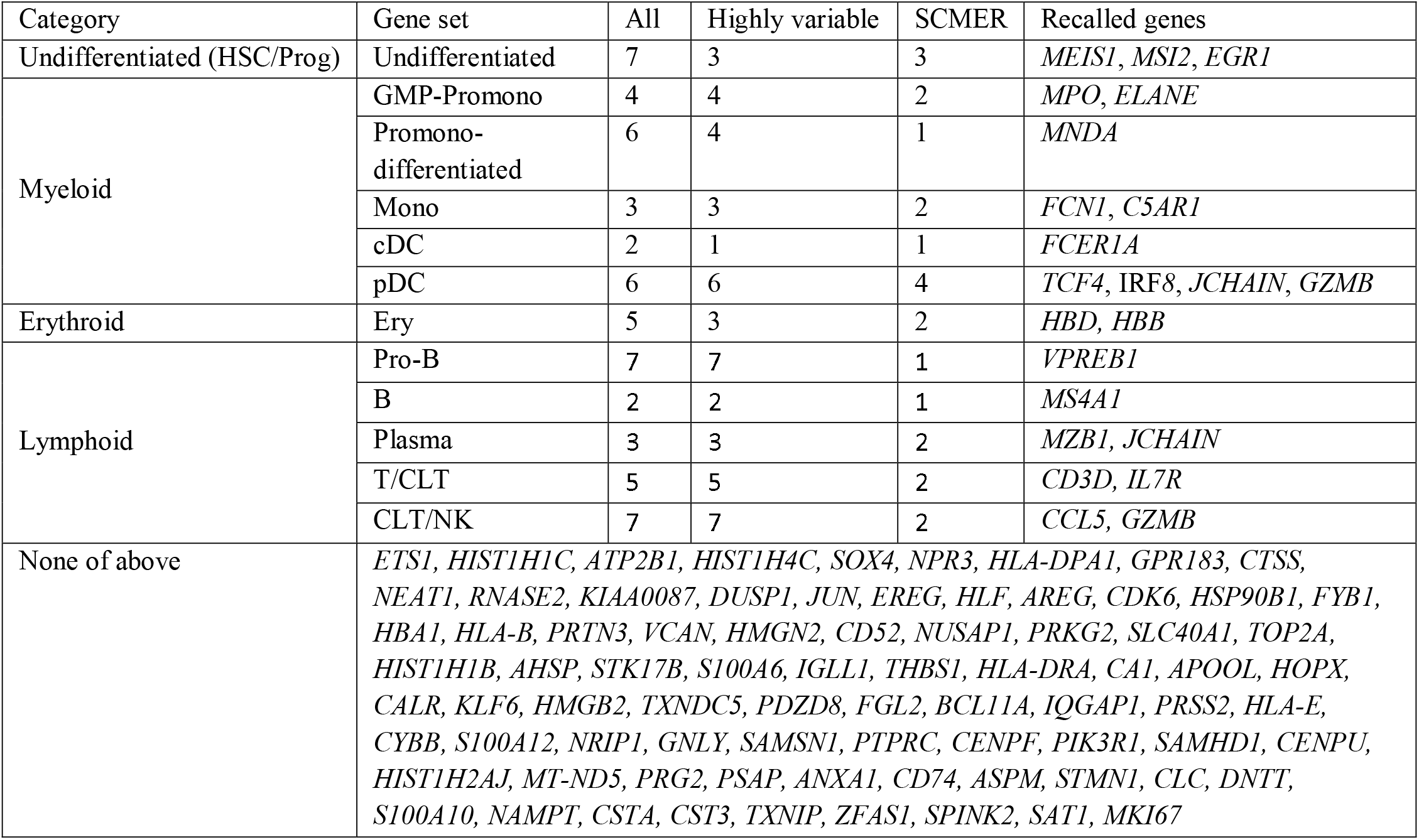
Bone Marrow features Recalled by SCMER

SCMER identified features delineating both clusters and sub-clusters. For example, the well-known lineage features such as *CD79A* (B cells) and *CD7* (T cells) and immune subtype markers such as *FCER2* (naïve B cells) and *ANKRD28* (TRM) were identified (**Fig. 3c,d** and **Supplementary Figure 4a**). Less reported features such as *SEPP1* for M2 macrophages were also among the list (**Supplementary Figure 4b**). The selected features also included genes that encodes lysozyme (*LYZ*), complements (*C1QA, C1QB*, and *C1QC*), granulysin (*GLNY*), and granzymes (*GZMA, GZMB, GZMK*, and *GZMH*) (**Fig. 3c and Supplementary Figure 4c**).

**Fig 4:**
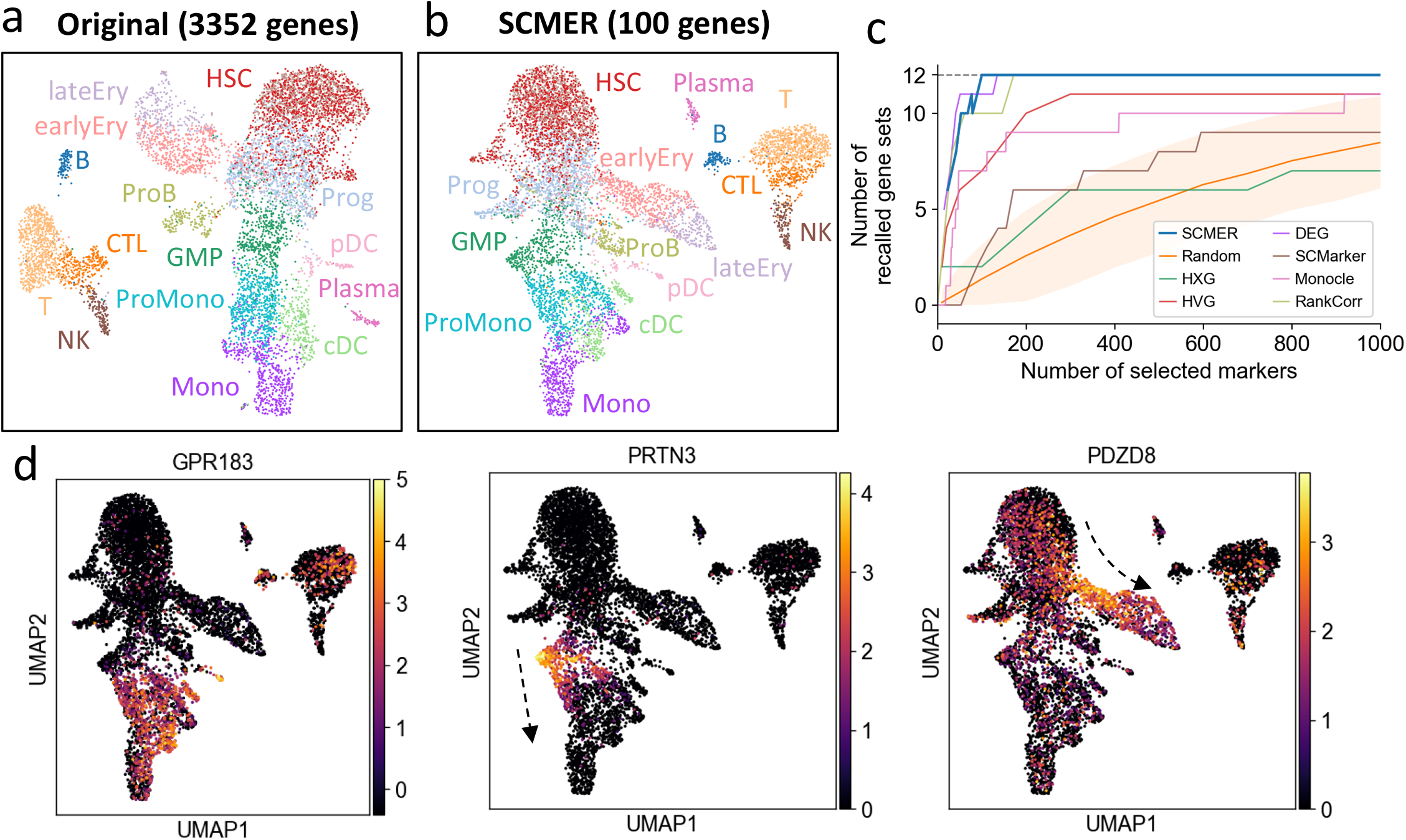
Results of the bone marrow hematopoiesis data. **(a)** UMAP embedding of the original dataset (List of abbreviations: **Supplementary Table 14**). **(b)** UMAP embedding of the dataset on SCMER selected genes. **(c)** Recall of gene sets for SCMER, scMarker, Monocle, HXG, HVG, PCA, and DEG, similar to **Fig. 2c.** **(d)** Activity of selected markers. Arrows are drawn for a visual reference for the developmental processes.

NK and ILC1 cells were mixed together in one cluster and can hardly be further dissected based on unsupervised clustering and DE analysis. However, based on the genes selected by SCMER such as *GNLY CCL4*, etc., which displayed dichotomizing levels within the cluster, we were able to further separate NK and ILC1 cells and estimate their abundances (**Supplementary Figure 5**).

**Fig 5:**
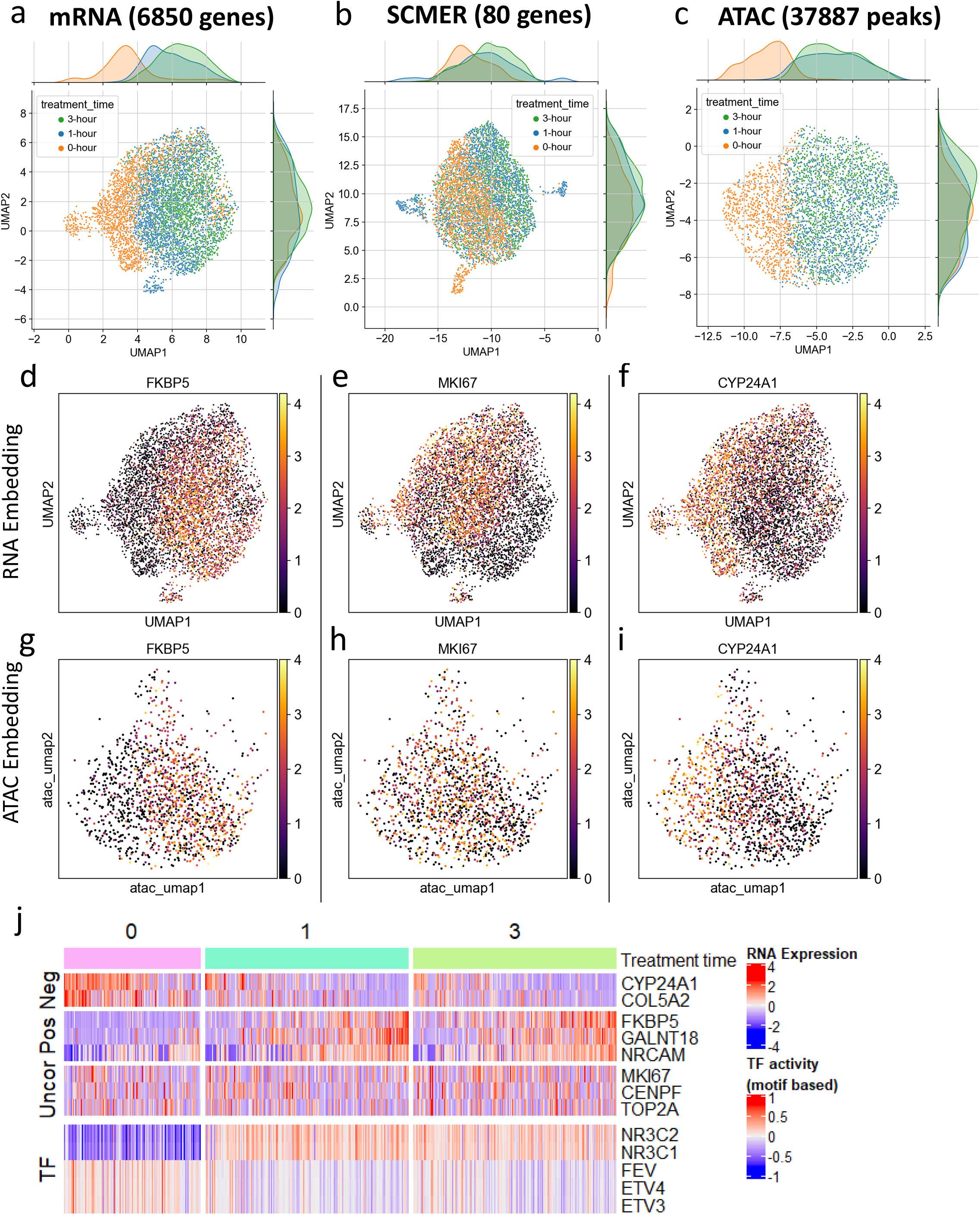
Results of the A549 lung cancer cell line data. (**a**-**c**) UMAP embedding of (a) the original sci-RNA-seq dataset, (b) the sci-RNA-seq dataset on SCMER selected markers, and (c) the sci-ATAC-seq peak dataset. (**d**-**i**) Expression of selected genes show in (d-f) RNA space and (g-i) ATAC space. ATAC space only includes co-assayed cells. (**j**) Heatmap of expression of selected genes and motif-based activity of highly variable transcription factors (TFs). (Uncor: uncorrelated, Pos: positively correlated, Neg: negatively correlated, with regard to *NR3C1* and *NR3C2*.) *ETV3* and *ETV4* are in the ETS transcription factor family.

SCMER also found TFs that regulate a wide range of cellular activities, including *JUN* and *FOS* (**Fig. 3d**), which are important for immune cell interactions. These features changed gradually among all the cell types, rather than expressing specifically in certain clusters. Other features such as *CD69* (known T cell activation feature) and *ODF2L* (novel T cell subtype feature) also showed gradual change among subtypes instead of clear-cut, on-and-off patterns (**Fig. 3f**). Notably, among our selected features that were not reported in the original publication, *DUSP1, DUSP2*, and *DUSP4* (**Fig. 3g and Supplementary Figure 4d**) were from a nucleus-predominant subfamily of dual-specificity phosphatases (DUSPs) family, which affect many cellular processes by regulating MAP kinases. *DUSP1* has been reported as a key regulator^45^ of both innate and adaptive immune responses by inactivating p38 and JNK (c-Jun N-terminal kinase). We found that *DUSP2* and *DUSP4* showed gradients majorly in the T cell cloud, while *DUSP1* was expressed in all the major cell types. These genes are related to a wide range of immune phenotypes including producing inflammatory cytokines and autoimmune responses, which are highly relevant to Crohn’s disease.

SCMER again compared favorably to the other methods that selected various numbers of features (**Fig. 3i, Supplementary Result 2**). It was evident that the other methods tended to ignore features associated with intra-cluster heterogeneity and multicellular programs. The novel genes selected by SCMER were also highly enriched in multiple immune pathways^43^ (humoral immune response, leukocyte migration, complement activation, etc.; **Supplementary Table 5**). Overall, SCMER sensitively preserved different types and levels of heterogeneity in the original data.

### 3.4 SCMER DISSECTS CONTINUOUS CELL LINEAGE DIFFERENTIATION

Cell differentiation involves many unique patterns of gene expressions, including those gradually changing, shared among cell types, or transiently activating during the process. None of these patterns can be characterized through clustering. Having shown that SCMER identifies both inter- and intra-cluster features, here, we examine if it can identify key differentiation features in hematopoiesis in human bone marrow (BM). A recent study^37^ sequenced 6,915 BM cells from four healthy donors. The transcriptomes of the cells formed a continuum instead of discrete clusters in the UMAP, reflecting the continuous differentiation process (**Fig. 4a**).

From this dataset, SCMER selected 100 features, which clearly preserved the trajectory of differentiation (**Fig. 4b**). The original publication reported 57 established features belonging to 12 sets (**Table 3**). SCMER recalled all the 12 sets as well as other well-known immune features (e.g. *GNLY* and *CD74*). SCMER also picked up features for less abundant cells that were not reported in the original publication (**Supplementary Table 6**), for example, *CLC* and *PRG2* for granulocyte and macrophage progenitor (GMP) cells (**Supplementary Figure 7a**). Neither of them was prioritized by DE analysis or Monocle^17^. Similarly, increasing *GPR183* expression was found in a subgroup of T cells, B cells, and Monocytes (**Fig. 4d**). The function of *GPR183* is not fully known, but emerging evidence shows that it may be a regulator of immune cell migration^46^.

SCMER identified many genes that displayed gradient along the developmental trajectories, for example, *AHSP* and *CA1* for Erythroid cells, and *VPREB1* for B cells (**Supplementary Figure 7b**). It also identified genes such as *PRTN3* (monocytes) and *PDZD8* (erythroid) that appeared transiently expressed during the developmental process and became dim in terminally differentiated cells (**Fig. 4d**), which were not prioritized by Monocle. Besides, it identified TF genes such as *JUN* and *SOX4*, which play important roles in regulating cell differentiation^47,48^. We comprehensively evaluated the performance and confirmed that SCMER outperformed the other unsupervised methods in recapitulating molecular diversity (**Fig. 4c**).

### 3.5 SCMER IDENTIFIES MOLECULAR DRIVERS FROM PERTURBED CELLS

More and more studies using single-cell technologies to investigate heterogeneity of cells in response to a genetic or chemical perturbation^49^. In these experiments, cell state may transition under complex kinetics.

To investigate the utility of SCMER in studying cellular responses, we applied it on single-cell data derived from dexamethasone (DEX) treated A549 lung adenocarcinoma cell line^38^. As reported in the original publication, the 1,429 cells sampled at 0, 1, and 3 hours after the DEX treatment formed a continuum in the transcriptomic space (**Fig. 5a**), indicating heterogeneous responses of the cell population. After running SCMER on the sci-RNA-seq data, 80 genes were selected, with the manifold and treatment states largely preserved (**Fig. 5b**).

We inferred TF activities based on motif enrichment^50^ in the chromatin accessibility (sci-ATAC-seq) data co-assayed on the same set of cells^38^ (**Methods, Fig. 5c**). Among the top 50 highly variable TFs (**Supplementary Figure 8a)**, *NR3C1*, the primary target of DEX^38^, had the most prominently increasing activity level over treatment time. Other TFs such as *FEV*^51^ and the ETS family^52^, also targets of DEX, had decreasing activity levels.

We then correlated the expression levels of the genes selected by SCMER with the activity levels of the top TFs. We found that *FKBP5, GALNT18, NRCAM*, etc. were positively correlated with *NR3C1*, while *CYP24A1, COL5A2*, etc. were negatively correlated (**Supplementary Table 7, Supplementary Figure 8**). In particular, *FKBP5*, a factor in the negative feedback loop of glucocorticoid receptor response and regulator of immune processes^53,54^, had the highest positive correlation (*r* =0.355) in the whole transcriptome; while *CYP24A1*, which regulates multiple metabolism processes^55^, was the most negative (*r* = – 0.365). Cells of high *FKBP5* expression levels came mostly from 1 and 3 hours (**Fig. 5d**), with matched polarized distributions in the RNA and the ATAC embeddings (**Fig. 5g**). Similar patterns were observed between cells of high and those of low *CYP24A1* expression levels (**Fig. 5f,i**).

Interestingly, SCMER also selected a group of genes uncorrelated with prominent TF activities (**Fig. 5j,Supplementary Figure 8**). Among them were *MKI67* (e.g., *r* = – 0.005with *NR3C1*) (**Fig. 5e,h**), which encodes proliferation marker protein Ki-67, and other cell-cycle genes such as *CENPF, TOP2A, RYBP, MLH3*, etc. Pathway analysis confirmed that these genes are highly enriched in cell proliferation pathways (**Supplementary Table 8**), indicating that an appreciable fraction of cells continued proliferating despite the treatment. It is not surprising that the levels of these genes were uncorrelated with chromatin state changes, as it has been shown that cell cycling status has little direct effect on chromatin accessibility^56^. Also among uncorrelated were several cancer cell stemness marker genes^43^ such as *ACTG1, TSC22D1*, and *FN1*, which may indicate that a fraction of cancer cells maintained their stemness during the course of the treatment. These genes would have been missed by a DE analysis supervised by the treatment time.

Taken together, our results demonstrated the superior power of SCMER in discovering features associated with heterogeneous cellular state change in the context of perturbation experiments.

### 3.6 SCMER ACHIEVES CROSS-MODALITY FEATURE MAPPING

One challenge in applying scRNA-seq for cell-typing is that expression levels of mRNAs can differ substantially from those of homologous proteins, due to post transcriptional modifications^57^. Although performing multi-omics assays may be the ultimate solution, they are currently associated with higher cost and lower throughput. Thus, rather than simply selecting the homologous mRNAs, it is beneficial to identify the set of genes whose expression levels maximally represent cellular diversity at the protein level. This capability can be important for designing targeted, cost-effective assays for preclinical and clinical applications. SCMER is ideally suited for such a purpose, as it allows selecting features in one modality while preserving manifold in another modality.

We ran SCMER on a CITE-seq dataset containing 14,468 bone marrow mononuclear cells (BMNC)^39^. The protein manifold based on 25 markers was utilized to “supervise” the selection of mRNAs (**Methods**). CITE-seq, which co-assays mRNA and protein markers from the same set of cells, is ideal for obtaining the optimal mapping between mRNAs and proteins (**Fig. 6a,b**).

**Fig 6:**
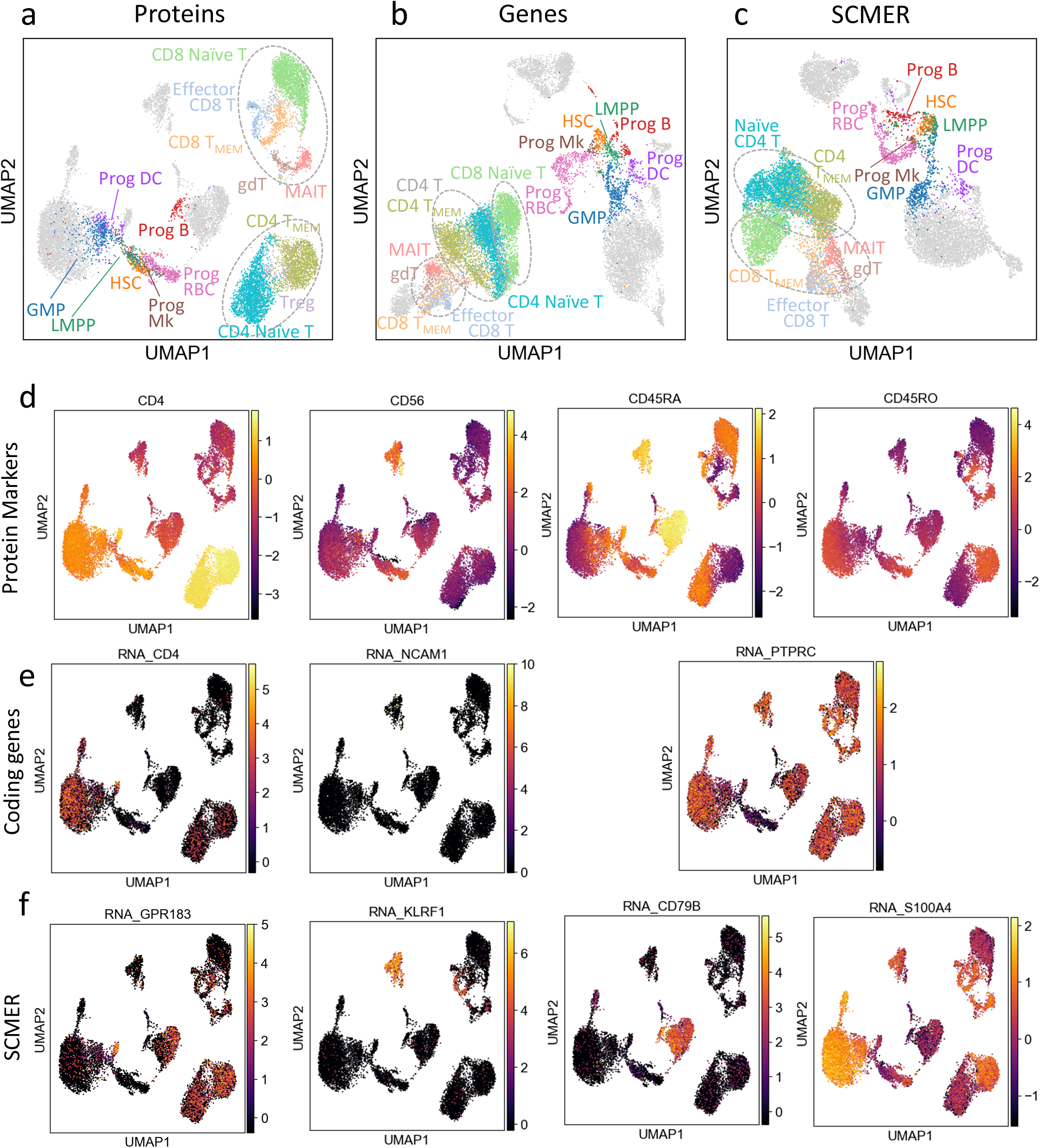
Results of the CITE-seq bone marrow mononuclear cells data. (**a**-**c**) UMAP embedding of original dataset using (a) protein, (b) genes, and (c) SCMER selected genes. T cells and Progenitor cells [HSC, LMPP (lymphoid-primed multipotent progenitors), GMP, and Progenitor of B, Mk (megakaryocyte), RBC, and DC cells], are highlighted for better visual identification. Fully annotated cell types are shown in **Supplementary Figure 10**. (**d**-**f**) Levels of representative (d) proteins, (e) genes, and (f) SCMER selected genes.

As shown, the mRNA expression levels of genes homologous to the protein markers, such as *CD4* (a T_h_ cell marker) and *NCAM1* (CD56, an NK cell marker) offered low power in delineating the corresponding cell types (**Fig. 6d,e**). Some markers, e.g., CD45RA (B cells and naïve T cells) and CD45RO (memory T cells) are isoforms of the same gene, *PTPRC*. Consequently, T cell subtypes were less distinguishable in the RNA space than in the protein space (**Fig. 6b**). The differences among CD8 T cell subtypes were even bigger than the differences between CD4 and CD8 T cells.

SCMER selected a set of genes that best preserved the diversity at the protein-level, notably the continuum among naïve CD8 T cells, memory CD8 T cells, and effector CD8 T cells (**Fig. 6c**). It identified genes that are non-homologous to the protein markers but better represent the protein level difference, for example, *GPR183, KLRF1, CD79B*, and *S100A4* for CD4, CD56, CD45RA, and CD45RO, respectively (**Fig 6d,f**). On the other hand, the SCMER result appeared to better delineate progenitor cells than the protein markers, which demonstrates a strength of integrating complementary modalities.

Similar conclusions were drawn when applying SCMER on another smaller PBMC CITE-seq dataset^40^ with 10 protein markers (**Supplementary Result 3**).

Importantly, the genes selected by SCMER from one donor (14,468 cells) appeared to preserve the cell diversity in another donor (16,204 cells) (**Supplementary Figure 10e-f**), which validated the applicability of SCMER in designing targeted panels for populational level testing.

## 4 Discussion

SCMER was designed to meet an important need in single-cell molecular data analysis, to sensitively identify non-redundant features that delineate both common cell lineages and rare cellular states ignored by current approaches. It provides an *ab initial* approach for discovering novel genes and features in high dimensional datasets, designing cost-effective assays for potential clinical applications, and assisting multi-modality integration of gene expression, proteins, and other features.

SCMER does not require clusters or trajectories and is not affected by uncertainties in clustering or trajectory inference. It explores alternative explanations via feature selection and reports the most salient features representing different facets of cells and underlying molecular activities. As a result, on datasets involving hematopoiesis, lymphogenesis, tumorigenesis, and drug resistance and response, SCMER identified features representing major cell types, rare cell populations (RCPs), continuous cell states, and multicellular programs. In our study, it prevailed existing unsupervised methods and often performed better than or comparably to the supervised methods when accurate labeling was possible (**Supplementary Figure 11**). Moreover, SCMER can handle batch effects by treating batches as a stratum, and finding a consensus set of features that preserve the manifold in respective batches (**Methods**). In that manner, it will prioritize genes contributing to biological but not technical variances.

SCMER can run in various supervised modes. It can accept a manifold from a different modality, for example, selecting RNA features under the guidance of a protein manifold (**Supplementary Result 3**). It can fix features preselected by users and find the best “partner” features (**Supplementary Result 4**) or select features from a shortlist (**Supplementary Result 5**). The framework appears effective on cell line and patient data generated by various technologies, including scRNA-seq and mass cytometry^41^ (**Supplementary Result 6**). This type of integrative analysis can potentially be extended to other modality combinations such as scRNA with scATAC, or mRNA with miRNA.

SCMER is an efficient method based on the orthant-wise limited memory quasi-Newton (OWL-QN) algorithm^31^. On a dataset with 10,000 cells and 2,000 candidate features, it typically converges in 20 to 40 iterations, which takes 5 to 10 minutes on a desktop computer equipped with a 3.20GHz 6-core Intel Core i7-8700 CPU. GPU acceleration is also supported, and the time consumption is halved with a middle-end NVidia GTX 960M GPU on a laptop computer with a 2.7GHz 4-core Intel Core i7-5700HQ CPU.

Because SCMER detects informative features that represent much wider and more complex biological processes than current methods, we expect it to be of immediate interest in projects producing large numbers of unsorted cells, such as the Human Cell Atlas^58^, the Human BioMolecular Atlas Program (HuBMAP)^59^, the Precancer Atlas^60^ and the Human Tumor Atlas Network^61^. It will be beneficial in various scenarios including biomarker discovery and clinical assay designing. As a first-of-its-kind method designed for manifold preserving feature selection on biomedical data, it can potentially be broadly applied to non-single-cell data, for example, bulk RNA expression^29^, copy number aberration^62^, and genetic and drug screening data in large cohort studies such as TCGA^63^, GTEx^64^, Depmap^65^, CTRP^66^, etc.

## 5 Methods

### 5.1 Cell-cell Similarity

SCMER is inspired by three methods: Stochastic Neighbor-Preserving Feature Selection (SNFS)^67^, t-distributed stochastic neighbor embedding (t-SNE)^27^ and Uniform Manifold Approximation and Projection (UMAP)^28^.

t-SNE is one of the most widely used method for data embedding. For a dataset **X** ∈ ℝ^*n*×*D*^ with *n* cells and *D* features, the similarity of a cell *i* to another cell *j* is defined as

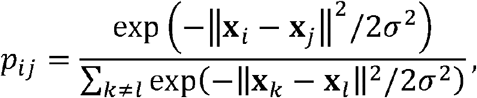

which comprises a cell-cell similarity matrix **P** ∈ ℝ^*n*×*n*^. *σ* is a scaling factor. It creates an *d*-dimensional embedding **Y**. ∈ ℝ^*n*×*d*^. It calculates another cell-cell similarity matrix **Q**. ∈ ℝ^*n*×*n*^ for **Y**, whose entries are

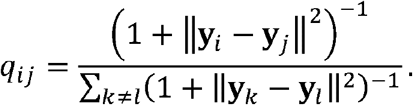

The cost function is defined as the Kullback-Leibler (KL) divergence of **P** and **Q**, formally

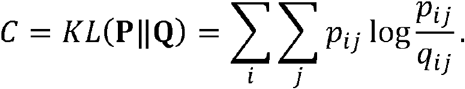

Because UMAP is more sensitivity to both global relationship between cell groups and local relationship within cell groups^29^, we borrowed a part of the UMAP formulation, i.e.,

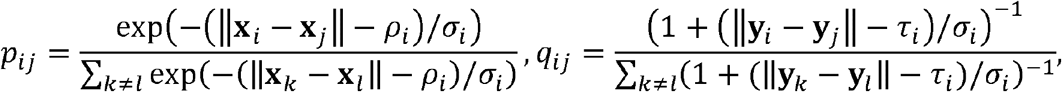

where *ρ*_*i*_ = min ║**x**_*i*_ – **x**_*j*_ ║ and τ_*i*_ = min ║ **y**_*i*-_ **y**_*j*_║ The scaling factor *σ*_*i*_ is chosen such that ∑_*j*_ exp (– (║ **x**_*i*_ – **x**_*j*_ ║ – *ρ*_*i*_) /*σ*_*i*_) = log_2_ *k*, which may be viewed as constructing a soft nearest neighbor graph. We default it to 100 in our experiments. Similar to UMAP, setting it in the range 10 to 1,000 gives very similar results^28^.

### 5.2 Marker Selection BY Elastic Net

Different from t-SNE and UMAP, instead of allowing **Y** to be an arbitrary matrix, we require each column of **Y** to be directly taken from a column of **X**, i.e., to select a feature. To formally model this procedure. We use a vector **W** ∈ ℝ^*D*^ to indicate the selection of the features, where 0 means unselected, and set

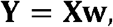

which set all unselected features to zero in **Y**. In terms of calculating the distances, zeroing out the columns is effectively discarding them. Thus, the definition of **Q** is unchanged. Ideally, to select *d* features, we optimize

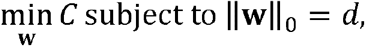

where ║**W**║ _0_ is the *l*_0_-pseudo-norm, i.e., the number of nonzero entries. However, this question is known to be NP-hard, whose determination requires checking all the 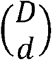 possibilities. Thus, we fall back to *l*_1_-norm, the convex approximation of *l*_0_ -pseudo-norm, as in

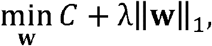

where *l*_1_-norm ║**w**║_1_ = ∑ _*i*_|w_*i*_ |and *λ* is the strength of the regularization. We denote the loss function as *L*. The number of chosen features decreases with larger *λ*. Thus, for a given *d*, we use a binary search to find a *λ*. Due to limitations of precision, the specific *d* may not always be achievable. In that case, we allow for a few more features to be selected, and discard those that are assigned with the lowest weights (Supplementary Note 2). In the result, the features who have nonzero weights in **w** are considered selected. The specific weight is not used in downstream analysis (**Supplementary Note 2**).

The cost, *C* = *KL*(**P**║**Q**),, is a robust indicator of whether the manifold is successfully retained. A typical range of *C* is 2.0 – 4.0 when the manifold is reasonably retained. More features (i.e., smaller *l*_1_-regularization) may be needed if the *C* is greater than 4.0.

Our model also allows an additional *l*_2_ -regularization (ridge) to form an elastic net model. It may improve the robustness of the panel by slightly increase the redundancy, so that noise or drop-out in one feature has less effects.

### 5.3 Batch Effect Correction BY Stratification

Batch effect is a common problem in experiments including multiple samples. For SCMER, the samples are considered a stratum. In specific, a set of **P** and **Q** can be constructed for each sample, denoted as **P**^(*i*)^ Batch effect is a common problem in experiments including multiple samples. For SCMER, the samples and **Q**^(*i*)^, while **w** is shared by all samples. A cost *C*^(*i*)^ can thus be calculated for each sample, and collectively form a new objective *C* = ∑ _*i*_ *C*^(*i*)^. Thus, SCMER will ignore features that identify different samples and focuses on features that retain cell-cell similarities in all/most samples.

### 5.4 Supervised Multi-omics Mode

To transfer the manifold in one matrix (**X**) to another (**X**′), either between different modalities or subsets of features of the same modality, we simply modify the definition of **Y** to **Y** = **X**′**w**. With all other procedures unchanged, the algorithm is now searching for features in **X**′ that gives a manifold similar to that of **X**. This is also applicable to select features from a shortlist of the original ones.

### 5.5 Using Preselected FEATURES

In the case that a researcher wants to specify a few features that are known to be useful, we slightly modify the regularization to *λ* ║**Vw**║_1_, where **V** = diag (**v**) is a diagonal matrix. If a feature is considered In the case that a researcher wants to specify a few features that are known to be useful, we slightly important *a priori*, the corresponding entry in **v** is set to 0 to avoid *l*_1_-regularization. In this “softly-supervised” way, SCMER is more likely to select these features, but may still discard some of them if they are contradicting with the manifold. Thus, in addition, we provide a “hard-supervised” way where a set of features are guaranteed to be kept. Other features are selected to supplement them.

### 5.6 Orthant-Wise Limited Memory Quasi-newton Algorithm

Limited-memory BFGS (L-BFGS) is an widely-used optimization algorithm in the quasi-Newton methods family^68^. It approximates the Broyden–Fletcher–Goldfarb–Shanno (BFGS) algorithm with *0*(*mD*) memory, where *m* can be chosen based on computing resources.

Although L-BFGS usually converge very fast (<20 iterations) for most *l*_2_ -regularized regression problems, it will diverge for *l*_1_-regularization, whose partial derivative is undefined at {**w** │ *w*_*i*_ = 0 ∃*i*}:

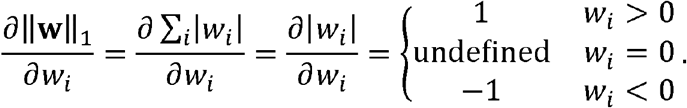

It should be noted that setting the undefined point to 0 (or any other value) at *w*_*i*_ = 0 does not solve the problem as the discontinuity will also break L-BFGS. A modified version of L-BFGS called orthant-wise limited memory quasi-Newton (OWL-QN) algorithm^31^ solves this problem by introducing pseudo-gradients and restrict the optimization to an orthant without discontinuities in the gradient.

L-BFGS optimizer is provided in PyTorch^30^, in which SCMER is implemented. Based on it, we implemented a special case of OWL-QN algorithm for optimization of the model. Two modifications we made are as follows.

Firstly, we derive the pseudo-gradient, where the pseudo-partial derivative at a discontinuity **w**_0_ of the loss function *L* = *C* (**w**) + *λ* ║**w** ║_1_ is defined as

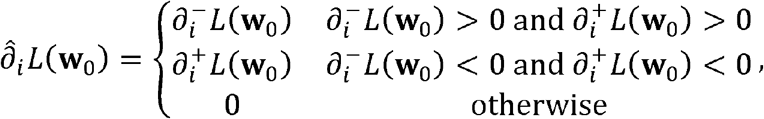

where 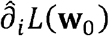 is the pseudo partial derivative and 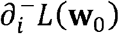 is the short hand of 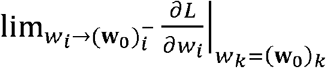, i.e, the left limit of the partial derivative. Similarly, 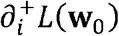 is the right limit.

Note that the gradient of *C*(**w**) is continuous, i.e., 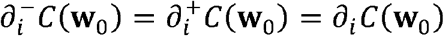. Thus,

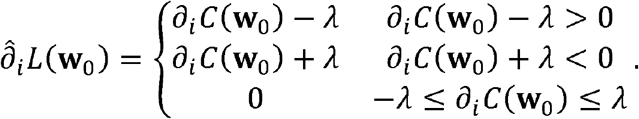

In fact, for *L*, discontinuities are {**w** │ *w*_*i*_ = 0 ∃ *i*}.

Secondly, we confine the search area in each quasi-Newton optimization step so that it does not cross any discontinuity. Specifically, for our problem where all discontinuity. Specifically, for our problem where all discontinuities are at 0, when updating **w**^*t*^ to **w**^*t*+1^, we reset the value of 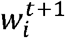 to 0 if 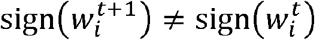. It constrains the optimization to be in the same “orthant” in each iteration.

### 5.7 Data Preprocessing

For the melanoma data^34^, which is TPM based, after removing ERCC spike-ins, we processed the data using the standard workflow of SCANPY^69^, including quality control (filtering out genes that are detected in less than 3 cells), normalization (10,000 reads per cell), log transformation, highly variable genes detection (with a loose threshold to filter out noisy genes; not to be confused with the DXG we compared with), and scaling.

For the Ileum Lamina Propria Immunocytes data^36^, bone marrow data^37^, and A549 data^38^, which are UMI based, we used the standard workflow of SCANPY, including quality control (filtering out genes that are detected in less than 3 cells), normalization (10,000 reads per cell), log transformation, highly variable genes detection, and scaling. We used the stratified approach to suppress batch effect on the Ileum Lamina Propria Immunocytes data.

For protein data in CITE-Seq^39,40^, we followed the preprocessing of protein data described in the original publication. For mRNA data in CITE-seq, we follow the standard workflow of SCANPY, as described above, except that we did not filter highly variable genes. We preprocessed protein data as mRNA data, without filtering highly variable genes.

### 5.8 Inference OF TF Activities

Because TFs tend to bind at sites with cognate motifs, accessibility at peaks with the motifs reflects their activity. To estimate transcription factor activity from sci-ATAC-seq data, we use chromVAR^50^ package with the default setting. It quantifies accessibility variation across single cells by aggregating accessible regions containing a specific TF motif. The observed accessibility of all peaks containing a TF motif is compared with a background set of peaks normalized for known technical confounders.

### 5.9 Comparison WITH Other Methods

To identify the highly expressed genes (HXG), we used the standard SCANPY^69^ workflow. HXG is defined by the total reads of a gene across all cells. To identify the highly variable genes, we followed the standard scoring method in SCANPY^69^.

SCMarker^19^ provides a gene list without ranks. It has two parameters, *n* and *k*, which affect the number of resulting features. Based on our observation, *n* has a minor effect on the result. Thus, we fixed *n* = 50 and tested *k* from 10 to 1,200to create feature gene lists of various sizes.

We ran Monocle^17^ in unsupervised and supervised manners. For the supervised run, the labels were used directly. The trajectory was inferred using clusters/labels and pseudo-time is calculated. Genes were ranked by the degree they are explained by functions (which were fitted with cubic splines) of pseudo-time. For the unsupervised run, we clustered the cells and visually confirmed the clusters are concordance with the labels.

We ran RankCorr^12^ in both supervised and unsupervised manner. For the supervised run, we used the label from the data directly. For the unsupervised run, we used the Leiden algorithm^70^ for clustering which is the recommended method in SCANPY. Default parameters were used, and the clusters are visually checked that they are reasonable.

For random results, we randomly selected gene sets of given sizes. Reported are mean performance and the critical level of statistically significantly better (or worse) than random as defined by single-sample one-sided z-test at 5% significance level.

## 6 Additional Information

### 6.1 Ethics approval and consent to participate

Not applicable in this study.

### 6.2 Consent for publication

Not applicable in this study.

### 6.3 Availability of data and material

The open source implementation of SCMER available at https://github.com/KChen-lab/SCMER under the MIT License. Scripts for reproducing all the results are also included. All original datasets are accessible through the original publications^34–41^.

### 6.4 Competing interests

The authors declare that they have no competing interests.

### 6.5 Funding

This project has been made possible in part by the Human Cell Atlas Seed Network Grant (CZF2019-002432 and CZF2019-02425) to KC from the Chan Zuckerberg Initiative DAF, an advised fund of Silicon Valley Community Foundation, grant RP180248 to KC and grant RP200520 to WP from Cancer Prevention & Research Institute of Texas, grant U01CA247760 to KC, grant U24CA211006 to LD, and the Cancer Center Support Grant P30 CA016672 to PP from the National Cancer Institute.

## 6.6 Acknowledgements

The authors would like to thank Hussein Abbas, Yuanxin Wang, Linghua Wang for their comments.

The authors acknowledge the support of the High Performance Computing for research facility at the University of Texas MD Anderson Cancer Center for providing computational resources that have contributed to the research results reported in this paper.

## 6.7 Authors’ contributions

SL, MM, WP, LD, and KC conceptualized the project. SL designed the SCMER algorithm and implemented the software. All authors collectively designed the experiments and analyzed the results. All authors drafted the manuscript. All authors have read and approved this paper.

## Notes

https://scmer.readthedocs.io/

### Competing Interest Statement

The authors have declared no competing interest.

