## Supplementary Text for "Single-Cell Manifold Preserving Feature Selection (SCMER)"

Supplementary Information for Single-Cell Manifold Preserving Feature Selection (SCMER)

Liang et al.

### 1 Supplementary Notes

#### 1.1 Supplementary Note 1: Illustrative Simulated Data

For **Fig. 1c**, we simulated 2000 cells for each branch. Each cell is randomly assigned a pseudo-time in $[0, 2]$, which is discretized into progenitor (time under 0.5), precursor (time between 0.5 and 1.5), and mature cells (time above 1.5). Expected values of features are determined as follows. Features in type I and II express on the corresponding cell types. Features in type III are monotonically correlated with time, on their corresponding branches. Features in group IV have a spike at time point 1.5. Expected values of noninformative features do not change. Dispersion was added by a negative-binomial distribution ($\theta=10$).

SCMER recalls all four types of markers in 45 markers. We find DEGs using Wilcoxon rank-sum test in a one-vs-all manner using the ground-truth cell type labels. It did not recall one type-II, III, or IV features in the first 100 features. The remaining features are mixed with noninformative features (noise). For pseudo-time-based correlation analysis, features with high linear correlation with the ground-truth pseudo-time are calculated. No type-IV feature is recalled in the first 120 markers. More advanced correlation analysis, such as cubic spline fitting, may be more powerful in finding these markers. The comparison is done on real data with Monocle.

To assess the cell type detection, we run $k$-nearest neighbor ($k$-NN) algorithm with $k=3$ in leave-one manner to evaluate the precision and recall of detecting each cell type using the selected markers. Specifically, for each cell, a classifier is trained on all cells except for itself. For each set of labels (major cell types / RCPs), each cell is either identified as one type, or rejected if no type wins the majority. With regard to a specific cell type, if the cells detected by the $k$-NN classifier is from the cell type, it is a true positive (TP). Otherwise, it is a false positive (FP). The precision and recall are defined as $\frac{TP}{TP+FP}$ and $\frac{TP}{TP+FN}$, respectively.

#### 1.2 Supplementary Note 2: Weights of Features Assigned by SCMER

Although SCMER generates rankings (weights) for the selected features, which represent the importance of the features to some extent, they should be interpreted with caution. If a smaller feature set is preferred, although it is acceptable to discard a few features with the least weights, we recommend rerunning SCMER with stronger $l1$-regularization in order to obtain a feature set of a different size. The reason is as follows. Given an optimal gene set of some size, if the desired size decreased by one, instead of only discarding one feature, it might be beneficial to also replace some of the other features in the current set. This is because of the nonlinear relations of features. For example, in a CyTOF PBMC dataset, if we currently have CD4 and CD8 representing CD4 T cells and CD8 T cells (together with other markers representing other cell types) and would like to further shrink the feature set by one, either discarding CD4 or CD8 may be suboptimal than discarding both and add CD3 for all T cells (provided that the feature set is too small to distinguish CD4 and CD8 T cells). Fortunately, the cost of rerunning SCMER is minimal given the efficiency of the algorithm.

Besides, our post feature-selection UMAP is not based on the weights. In other words, all nonzero weights are considered one regardless of the specific values.

### 2 Supplementary Results

#### 2.1 Supplementary Result 1: SCMER Captures Intra-Cluster variation in 198 cancer cell-lines and patient tumors

To further verify that SCMER is sensitive to features that drive intra-cluster heterogeneity and discriminate RCPs, we applied SCMER on a recent pan-cancer study pooling 198 cancer cell lines from 22 cancer types^1^. The original study identified 12 recurrent heterogeneous programs (RHPs) from multiple cell lines in the categories of cell cycle, senescence, stress and interferon responses, EMT, and protein metabolism. Also provided were 21 RHPs found in patient tumor samples. Individual cell lines were well separated out in the UMAP (**Supplementary Figure 3a**) while RHPs contributed mostly to within-cell-line (cluster) heterogeneity. Consequently, the top DE genes of each cell line (173 in total) did not include marker genes in the 12 RHPs such as the G1/S and G2/M phases of cell cycle, protein maturation, or proteasomal degradation (**Supplementary Table 3**). In contrast, SCMER was able to recall all but stress response with the same number of features. SCMER identified novel genes such as *PRDX1* (proteasomal degradation marker) and *CDC20* (Cell cycle G2/M marker), which were clearly RHPs in most cell lines (**Supplementary Figure 3b**) but missed by the DE analysis. SCMER also recalled genes in all the reported 21 RHPs derived from patient tumor samples, while DE analysis failed to recall stress response markers derived from head and neck squamous cell carcinoma (HNSCC), and markers for G1/S and G2/M cell-cycle phases. Clearly, SCMER is sensitive to intra-cluster heterogeneity even in the presence of hundreds of clusters.

#### 2.2 Supplementary Result 2: Assessment of SCMER on Immunocytes

To perform an objective comparison on the Ileum lamina propria immunocytes dataset, we curated 31 gene sets from the original publication (**Supplementary Table 4**). Except for two that had no highly variable genes (i.e., not possible to recall because SCMER was run on highly variable genes), SCMER recalled 23/29 based on 250 features. Although features were not recalled for some cell types, those cell types were still clearly present in the UMAP (**Fig. 3b**). In fact, alternative features for those cells were found by SCMER. For example, *MAF* is a specific feature for Lymphatic cells that dictates its differentiation^2^ (**Supplementary Figure 4e**). Other examples included *FCER1G* for DC2^3^, *ID2* for DC1^4^, *ITGA1* for glial cells. Some cells could be recognized by using multiple features, for example *COL1A2*+ and *CCL11*- specifically identified smooth muscle cells (**Supplementary Figure 4e**). SCMER also found novel features including *TFF3* (lymphatic cells), *CYB561A3* (B cell subtypes; a paralog of *CYBD1*), and *ODF2L* (T cell subtypes) that were not previously reported in the original paper, but clearly discriminated cell types or cell states (**Fig. 3f and Supplementary Figure 4e**). Because SCMER is designed to select non-redundant features, these alternative features may preclude it from finding the established markers.

#### 2.3 Supplementary Result 3: SCMER Finds Features Transferring Manifold Between Modalities

Toward an intuition of the selected features, we experimented on a peripheral blood mononuclear cell (PBMC) dataset produced by CITE-seq^6^, which has matched proteomic and transcriptomic profiles for each cell. Although the proteome data only contains 10 surface features, the cells cluster better on it than the much larger transcriptomic data (**Supplementary Figure 12a**,**13a**), where CD4 T cells and CD8 T cells are intermingled. Using the 14 transcriptomic counterparts of the surface features also results in bad separation of cell types (**Supplementary Figure 13b**), because large discrepancy shows in the transcriptomic and proteomic levels (e.g. CD4 vs *CD4*, CD11c vs *ITGAX*, CD45RA vs *PTPRC*, and CD57 vs *B3GAT1*; **Supplementary** **Figure 12c,14**). In fact, many of the corresponding genes are not even in the top ten most correlated ones with the features (**Supplementary Table 9**). Even for the features that are highly correlated with their mRNA counterpart, other gene may appear to be even higher correlated (e.g., CD2 vs *HLA-DRA*, CD14 vs *LYZ*, and CD19 vs *CD79A*).

We used SCMER to also keep 14 genes. Using these features, the embedding of cells (**Supplementary Figure 12b**) has better separation of CD4 T cells, CD8 T cells, and NK cells compared with the embedding on all genes (**Supplementary Figure 13a**), the corresponding genes of the surface features (**Supplementary Figure 13b**), and genes that are highly correlated with the surface features (**Supplementary Figure 13c,d**). The selected features are clearly more correlated with the surface features compared with their transcriptomic counterparts (**Supplementary Figure 12d**). In fact, 13 out of 14 selected features (except for *FLT*) are in the top 10 correlated genes of the surface features (**Supplementary Table 9**). We noticed that no features are found for CD57 by SCMER. The reason might be that CD57 appears to contribute the least to the manifold (**Supplementary Figure 14j**). When we loosened the restriction of number of features to 26, the embedding further improves (**Supplementary Figure 12e**) and genes correlate with CD57 are also recalled (**Supplementary Figure 13f**, **Supplementary Table 10**). SCMER addresses the problem of discordant mRNA and protein levels and helps generate embedding/clusters similar to what is generated by surface feature profile using mRNA data. The whole transcriptome can still be used to identify cell type/states afterward. This type of analysis may also be applicable to scRNA and scATAC, or mRNA and miRNA.

#### 2.4 Supplementary Result 4: SCMER Finds Features Supplements a Preselected Feature List

In **Supplementary Results 3**, we showed that some genes are not the most correlated one of their protein counterparts. That said, when designing a panel, it may still be desirable to keep these genes for better biological interpretability. In light of this need, we did the same experiment on the data, but fixed the 14 genes correspond to the surface markers [*CD2, CD3D*, *CD3E*, *CD3G, CD4, CD8A, CD8B, ITGAX* (CD11c)*, CD14, FCGR3A* (CD16)*, FCGR3B* (CD16)*, CD19, PRPRC* (CD45RA), and *B3GAT1* (CD57)]. SCMER found 26 additional genes (**Supplementary Figure 15a** and **Supplementary Table 11**). SCMER reduced the number of features for the surface markers that are already well-characterized by the preselected features (e.g., CD3 and CD8), and added more features for those that are not (e.g., CD4, CD11c, CD19, CD49RA, and CD57). The manifold is well maintained by these markers (**Supplementary Figure 15b** and **Supplementary Figure 13**).

#### 2.5 Supplementary Result 5: SCMER Can Focus on Novel feature Discovery

In the experiments above, both established features and novel features are detected. SCMER can be set to discover only novel features. Having confirmed that SCMER transfers manifold information between modalities, we use the same strategy to better detect novel features. Just like choosing transcriptomic features to mimic the proteomic manifold, we choose features from genes that are not established features to mimic the original manifold defined by all genes. We demonstrate this application on the bone marrow dataset, where the established features are excluded from the candidate features (while they are still used to define the original manifold). Without using the most well-established feature genes, SCMER can still select a set of features to recover the manifold (**Supplementary Figure 15c** compared with **Fig. 4a**). The selected features concords well with cell types and developmental trajectories (continuums) (**Supplementary Table 12,13**), which shows that SCMER reliably discovers features from a less studied set of genes to inspire novel findings.

#### 2.6 Supplementary Result 6: CyTOF Data

We validated that SCMER is also applicable to CyTOF data, as the dimensionality differs from scRNA-seq. We used an AML dataset including 104,184 labeled cells^7^ with 32 protein markers (**Supplementary Figure 16a**). We used SCMER to find a subset of markers. Although the panel size is already small and most markers are essential for distinguishing the cell types, we were still able to squeeze it to 19 markers (**Supplementary Figure 16c**) while keeping the manifold (**Supplementary Figure 16b**). Although selecting markers from a specially designed CyTOF panel may not help suggest novel markers, combining the most representative combination of markers from a few cell type specific panels to form a comprehensive panel. As a comparison, we randomly chose 19 markers and calculated the UMAP embedding and repeated it five times (**Supplementary Figure 16d**). SCMER clearly preserves the manifold better.
