## Supplementary Figures for "Single-Cell Manifold Preserving Feature Selection (SCMER)"

### Supplementary Fiugre 1

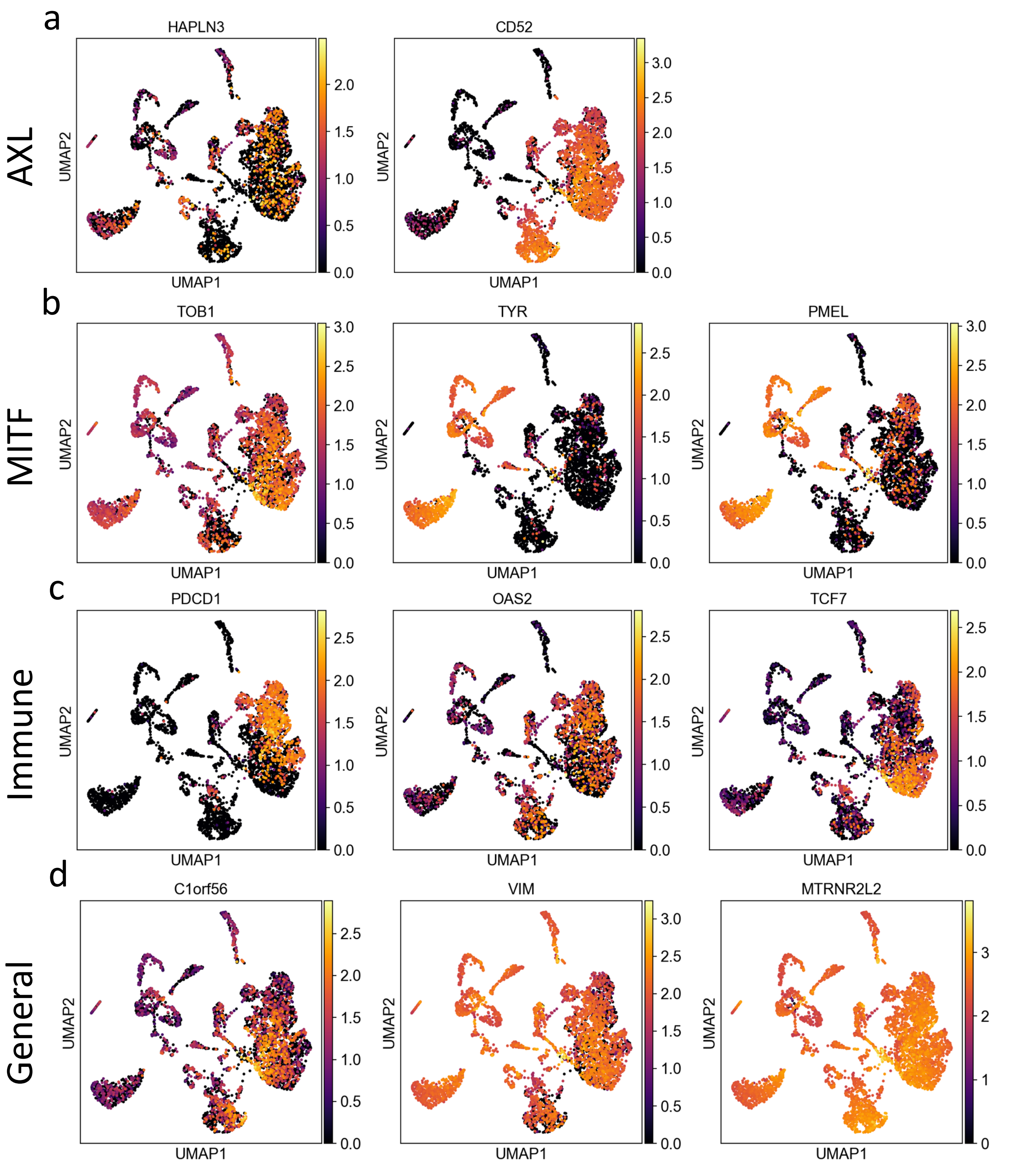


**Supplementary Figure 1 related to Fig. 2:** Additional results on the Melanoma data. (**a**-**d**) Examples of SCMER selected genes in (a) AXL and (b) MITF programs, which are intratumoral heterogeneous and related to resistance, and (c) immune cells and (d) all cell types. The markers listed here are not comprehensive.

Supplementary Fiugre 2

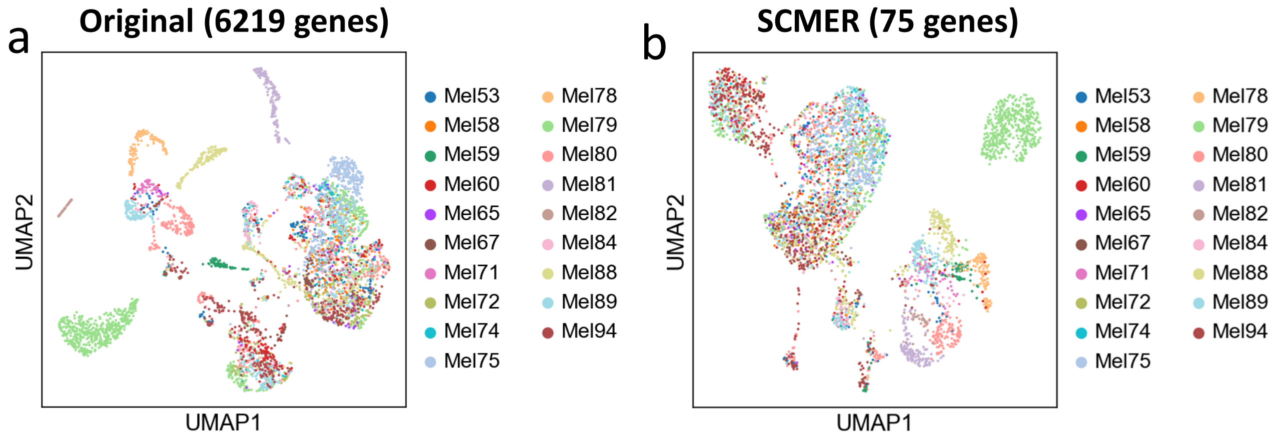


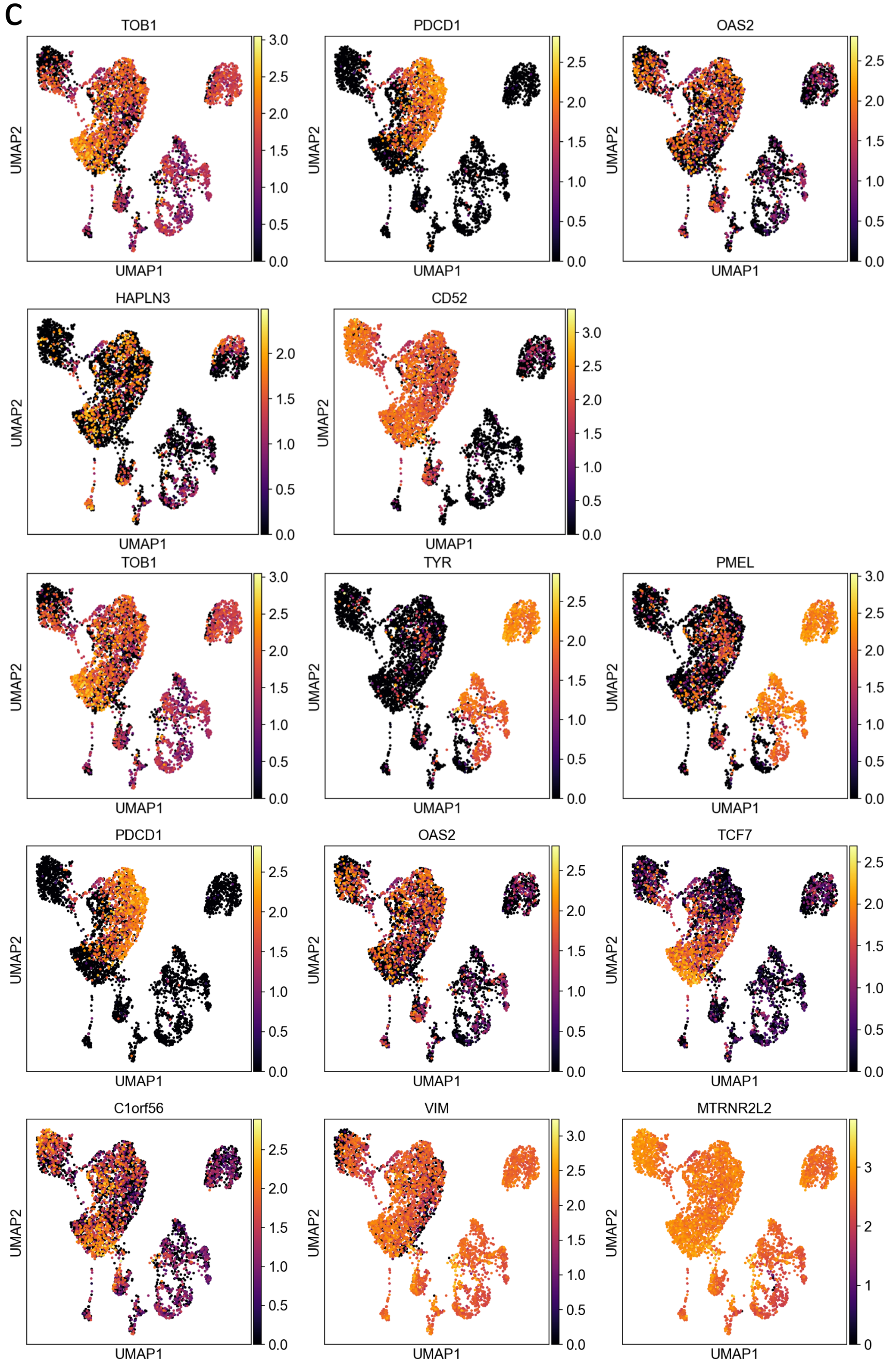


**Supplementary Figure 2 related to Fig. 2:** Additional results on the Melanoma data.
(**a**-**b**) UMAP embedding of the dataset (a) without and (b) with feature selection, colored by patients.
(**c**) All genes shown in **Fig. 1** and **Supplementary Figure 1** embedded in the manifold of genes selected by SCMER.

Supplementary Fiugre 3

a

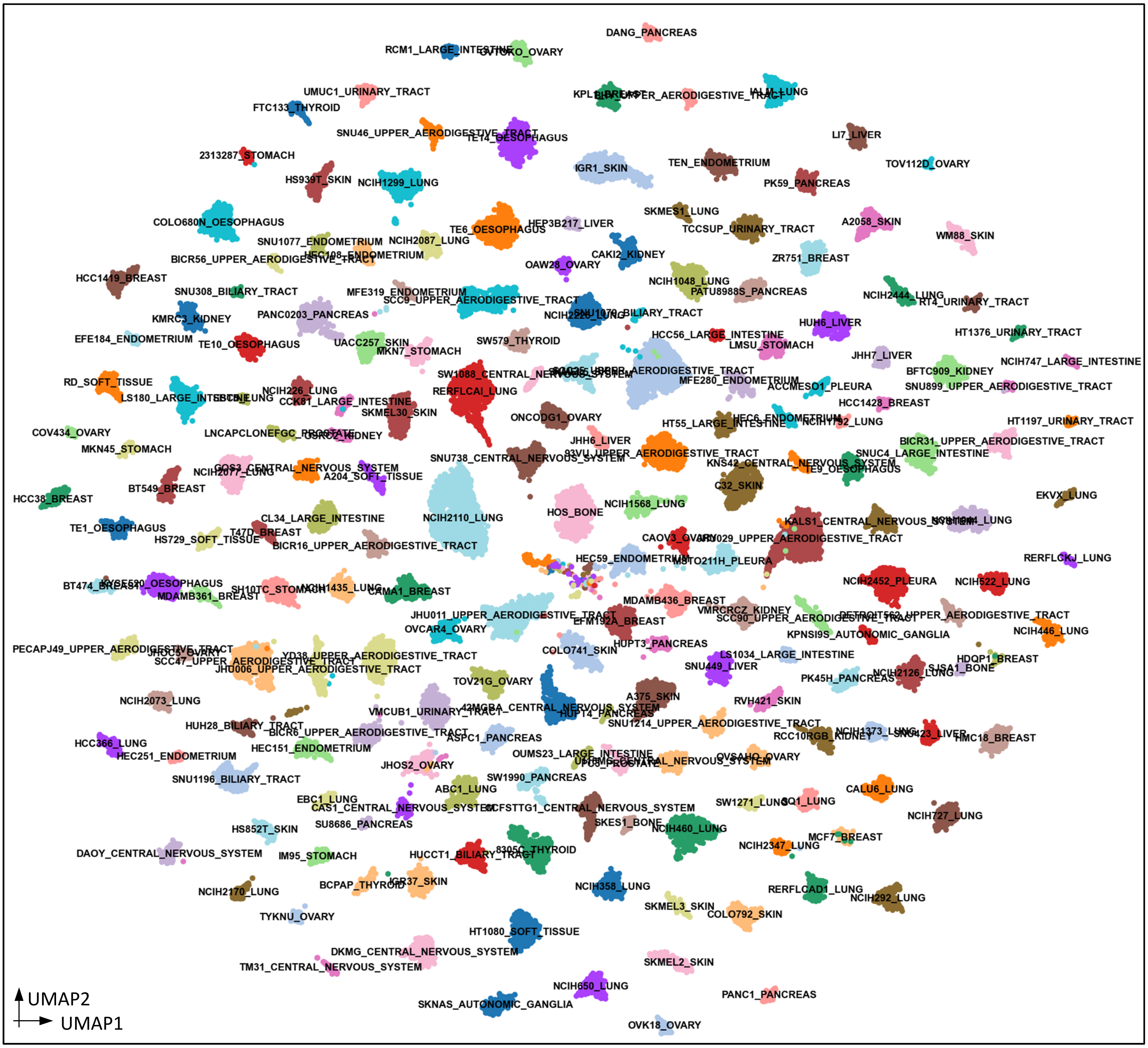


b

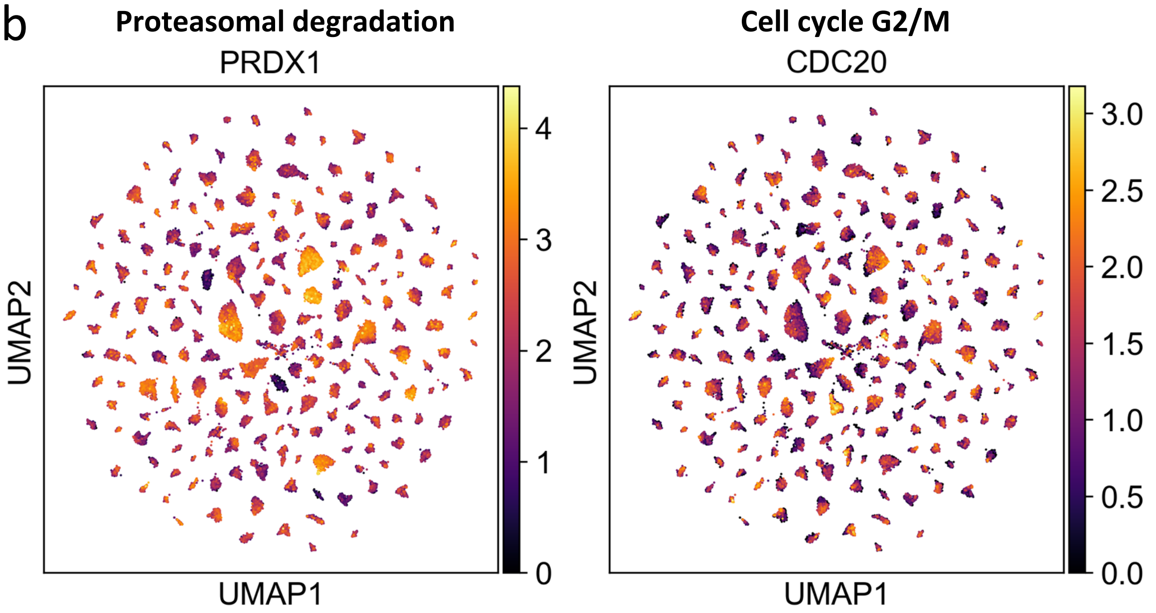


**Supplementary Figure 3:** Results of the pooled cancer cell line data. (a) UMAP embedding of the cancer cell lines. Due to the large number of cell lines, colors are reused and may be the same for multiple cell lines. (b) Expression of representative genes that reflect recurrent heterogeneous programs shared by a majority of cell lines.

Supplementary Fiugre 4

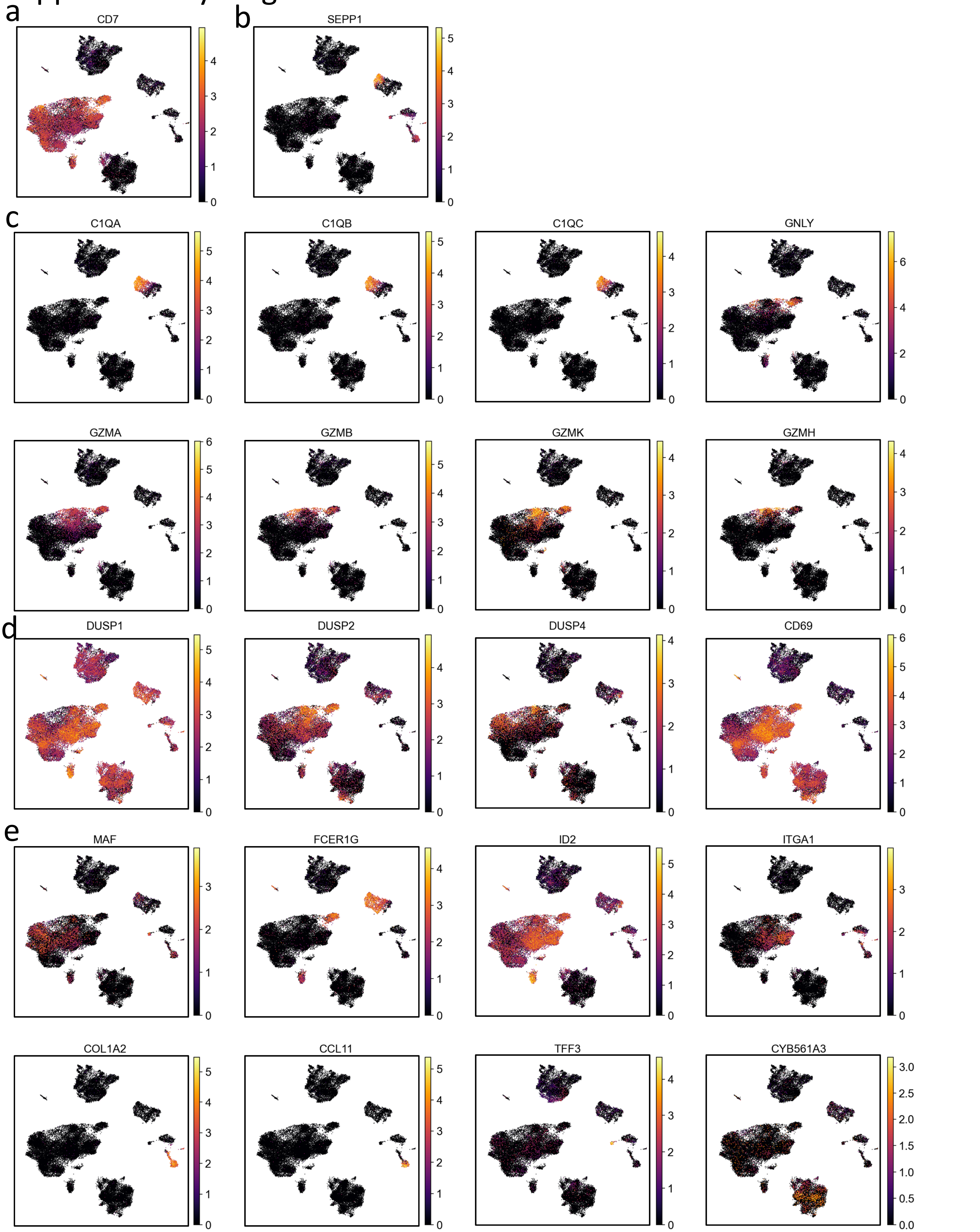


**Supplementary Figure 4 related to Fig. 3:** Expression of other markers mentioned in the main text for the ileum lamina propria immunocytes data.

### Supplementary Fiugre 5

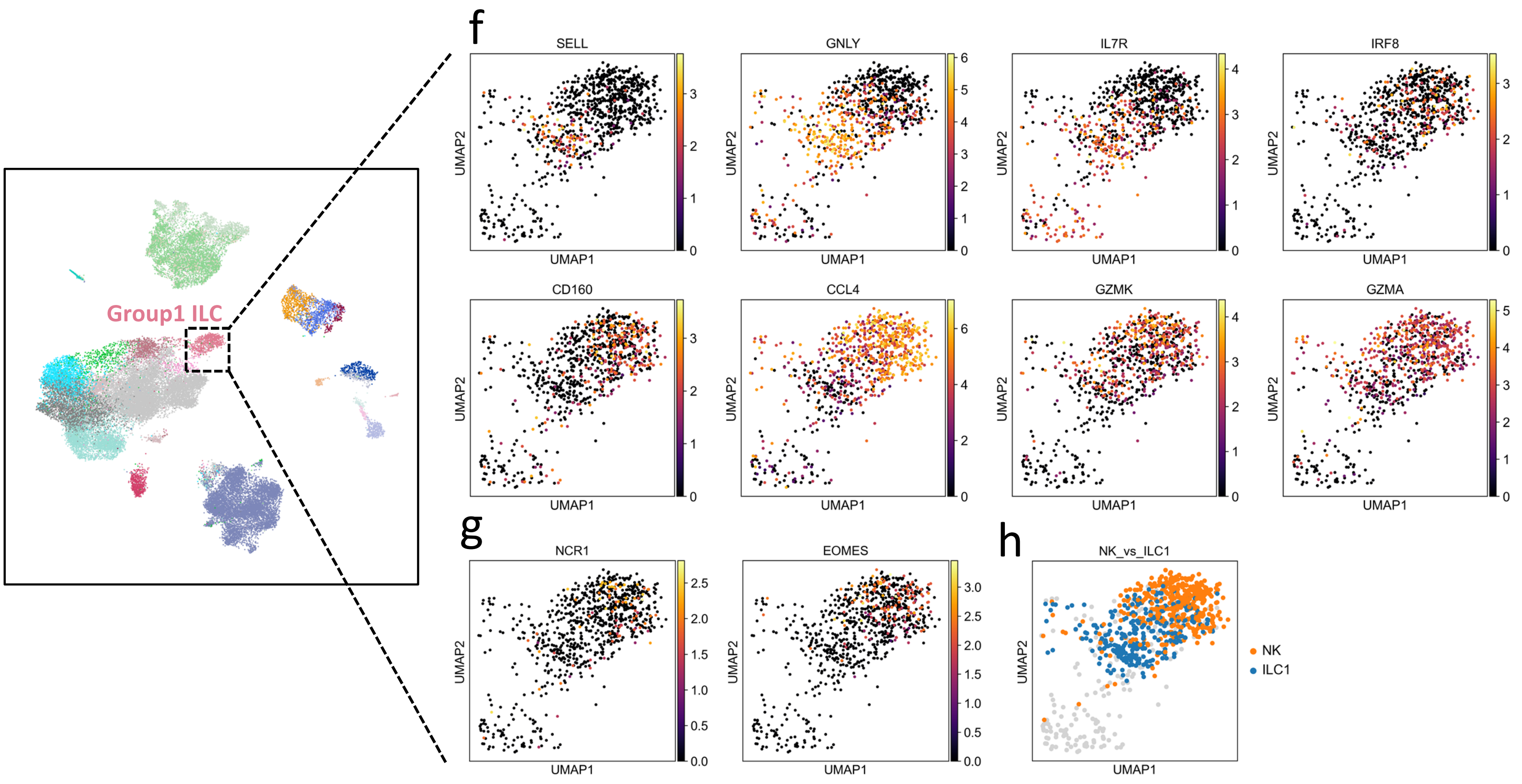


**Supplementary Figure 5 related to Fig. 3:** Separation of NK cell and ILC1.
(**a**) Location of Group 1 ILC (including NK cell and ILC1).
(**b**) Expression of markers separating NK cell from ILC1 selected by SCMER.
(**c**) Expression of other known markers for NK cells. They are less informative makers due to their sparsity, but nevertheless consistent with markers found by SCMER.
(**d**) NK (61.8%) and ILC1 (38.2%) cells inferred by their preferentially expression of genes selected by SCMER. Other cell types are colored in gray.

### Supplementary Fiugre 6

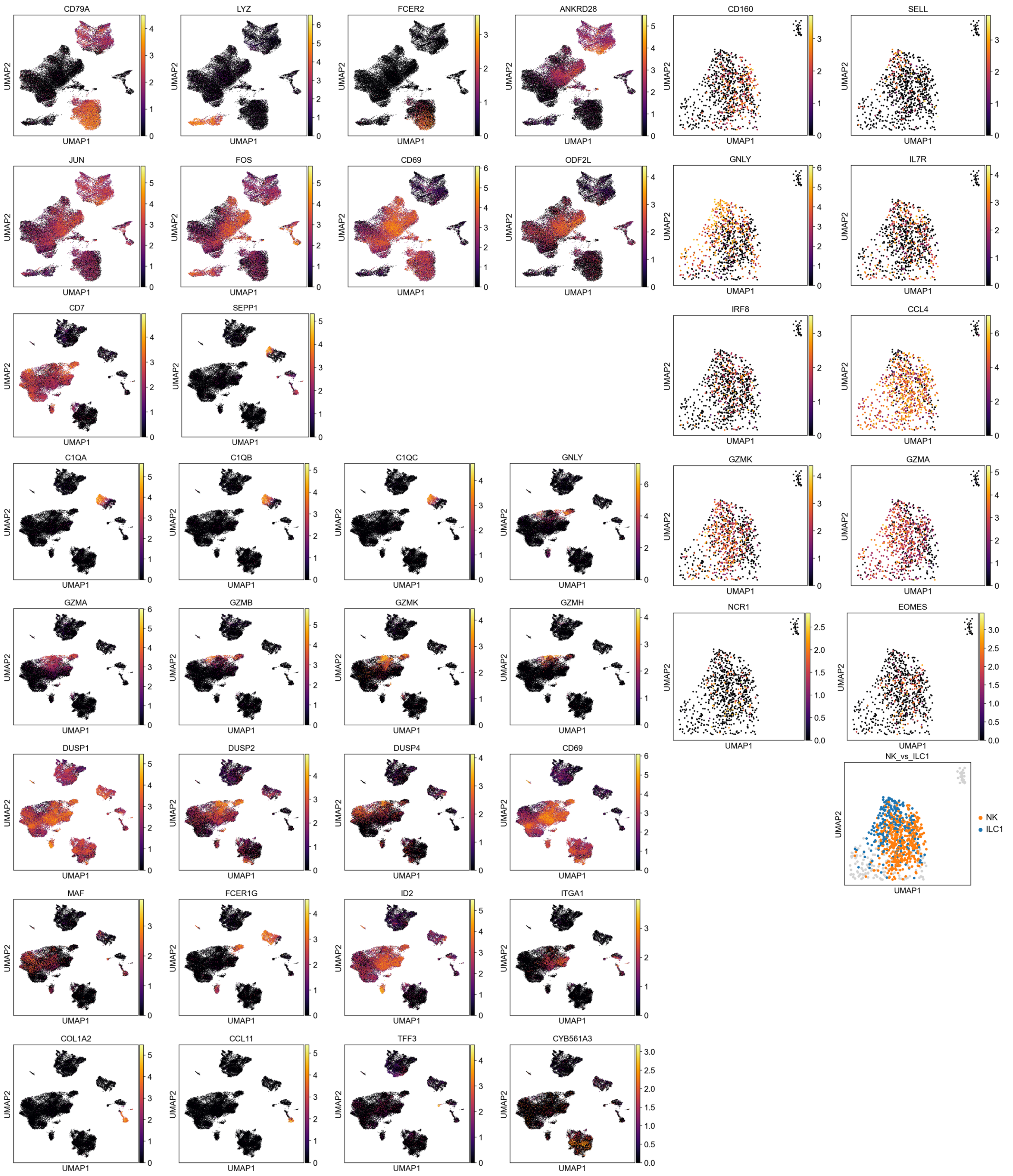


**Supplementary Figure 6:** Expression of all genes shown embedded in the manifold of genes selected by SCMER.

### Supplementary Fiugre 7

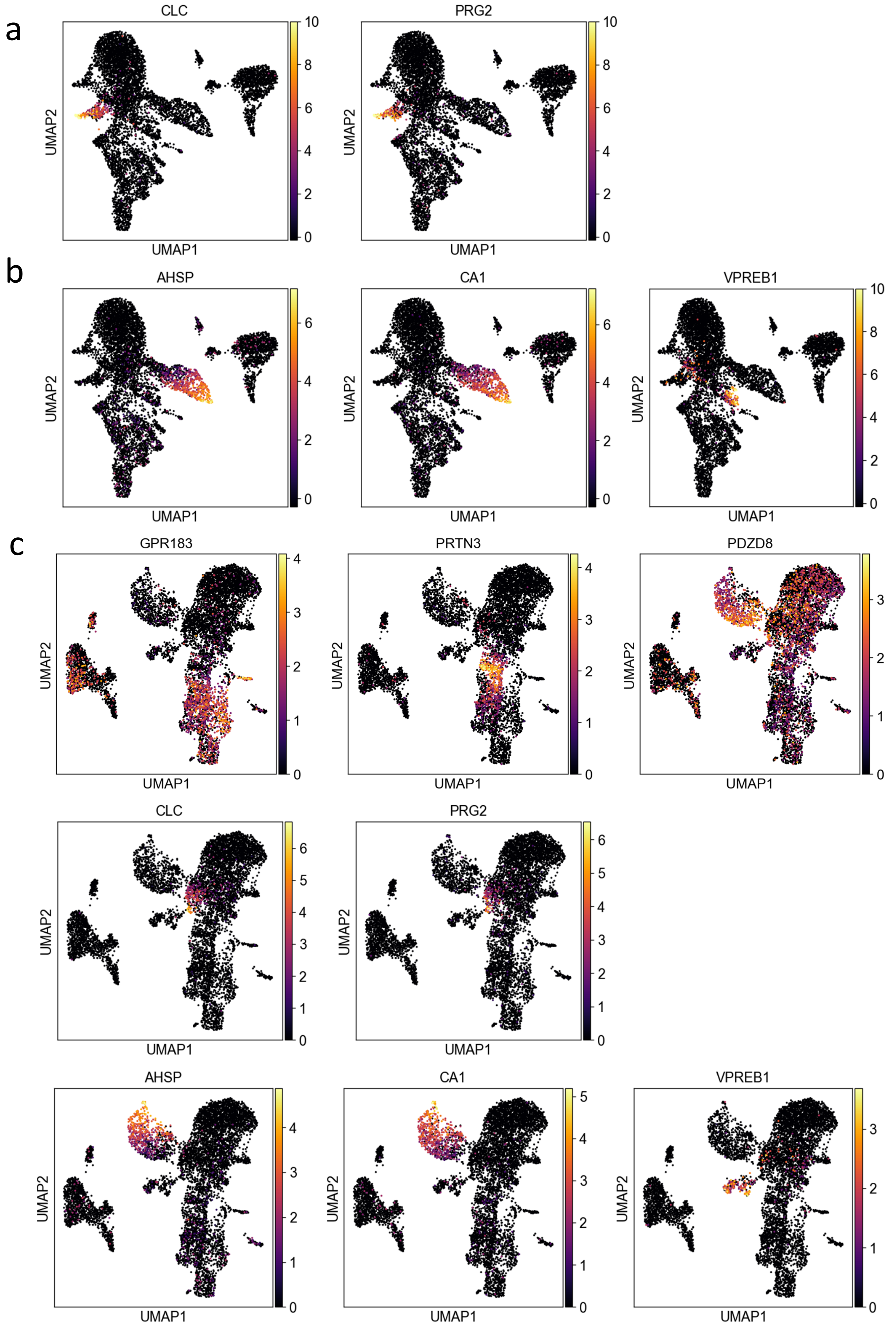


**Supplemental Figure 7 related to Fig. 4:** Additional results on the bone marrow hematopoiesis data.
(**a**,**b**) Activity of other selected markers mentioned in the main text.
(**c**) All genes shown above embedded in the original manifold.

Supplementary Fiugre 8

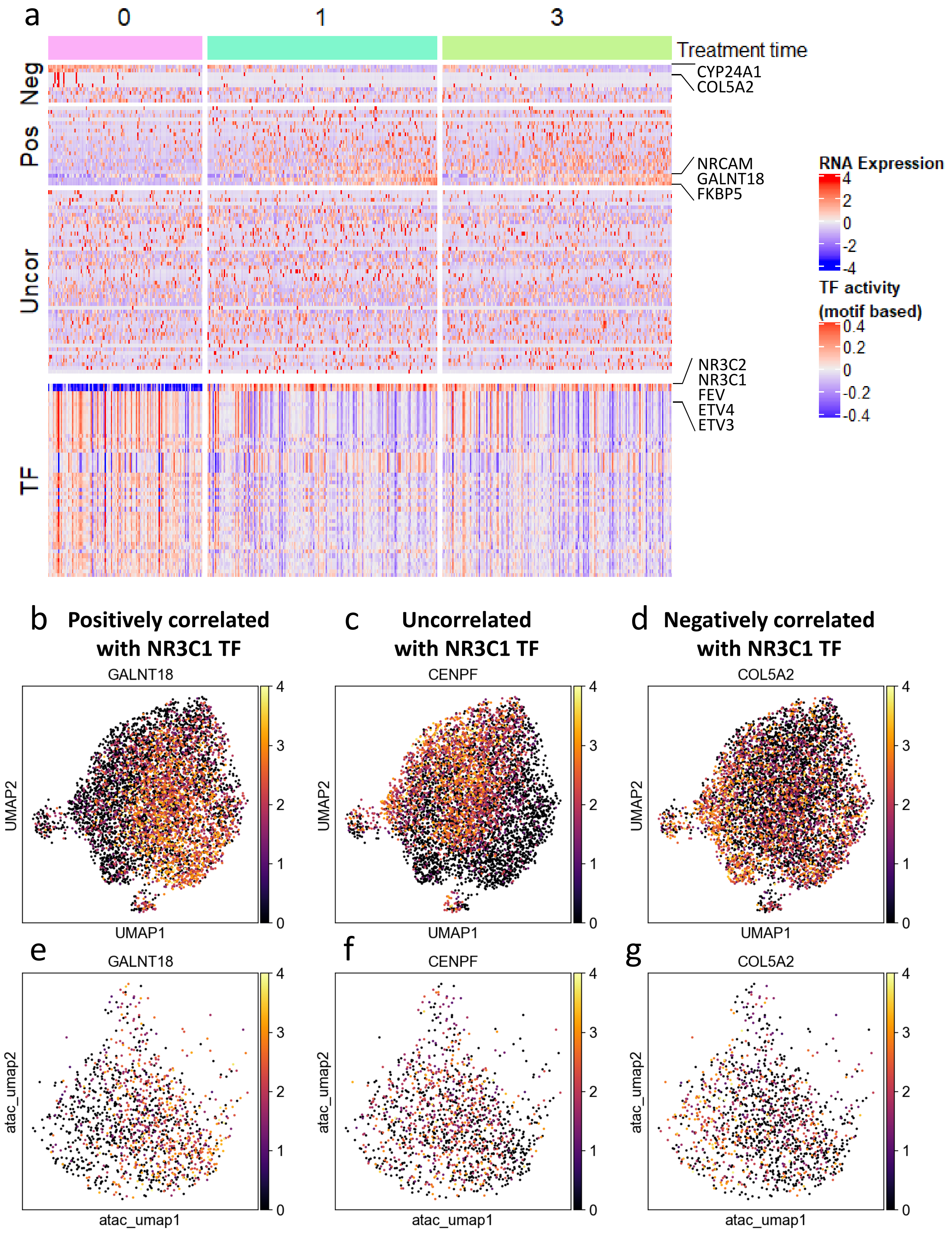


**Supplementary Figure 8 related to Fig. 5: Additional results on the A549 lung cancer cell line.**
(**a**) Heatmap of all genes selected by SCMER and 50 highly variable transcription factors (TFs).
(**b**-**g**) Expression of selected genes show in (b-d) RNA space and (e-g) ATAC space.

### Supplementary Fiugre 9

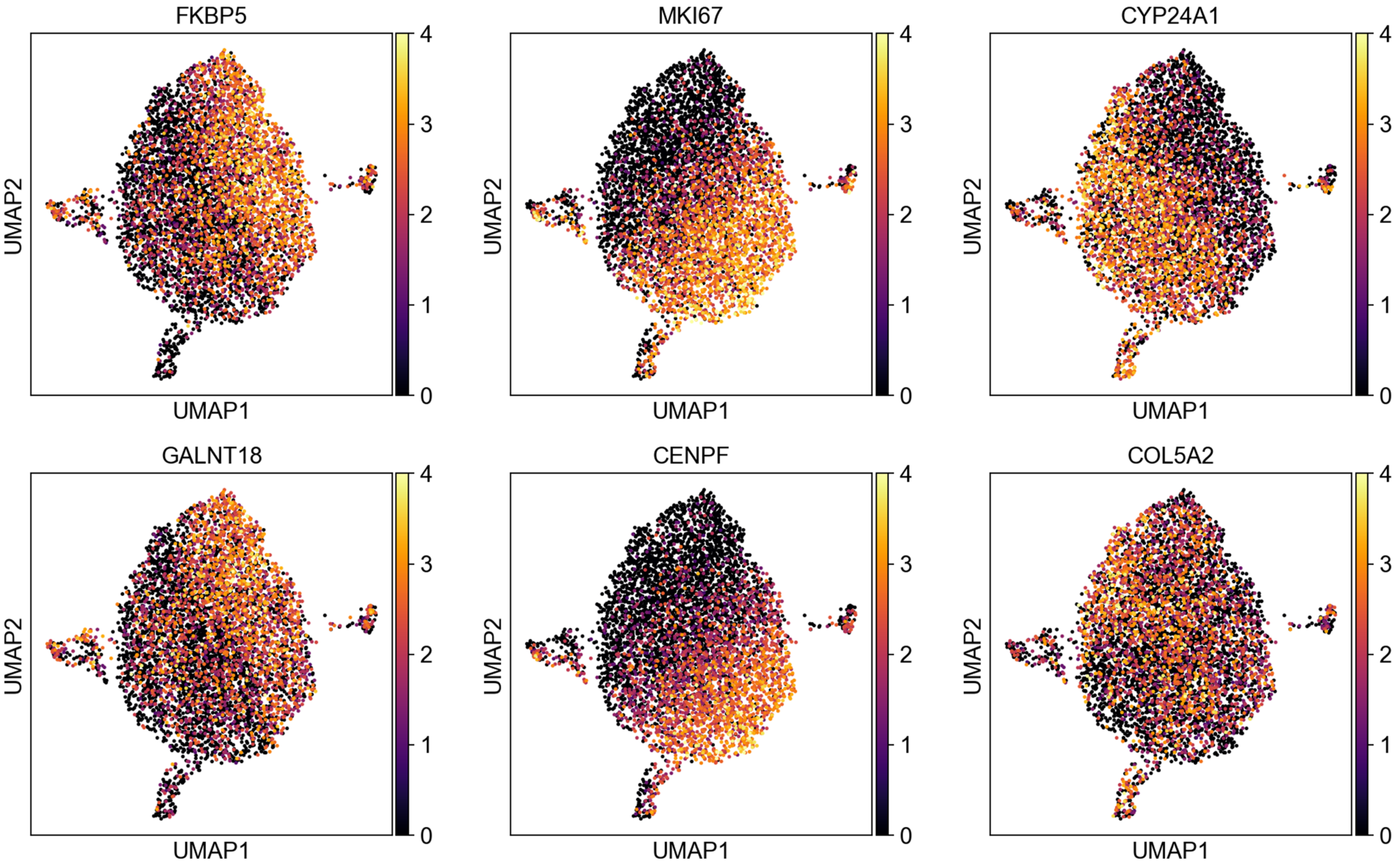


**Supplementary Figure 9 related to Fig. 5:** All genes shown embedded in the space of SCMER selected genes.

Supplementary Fiugre 10

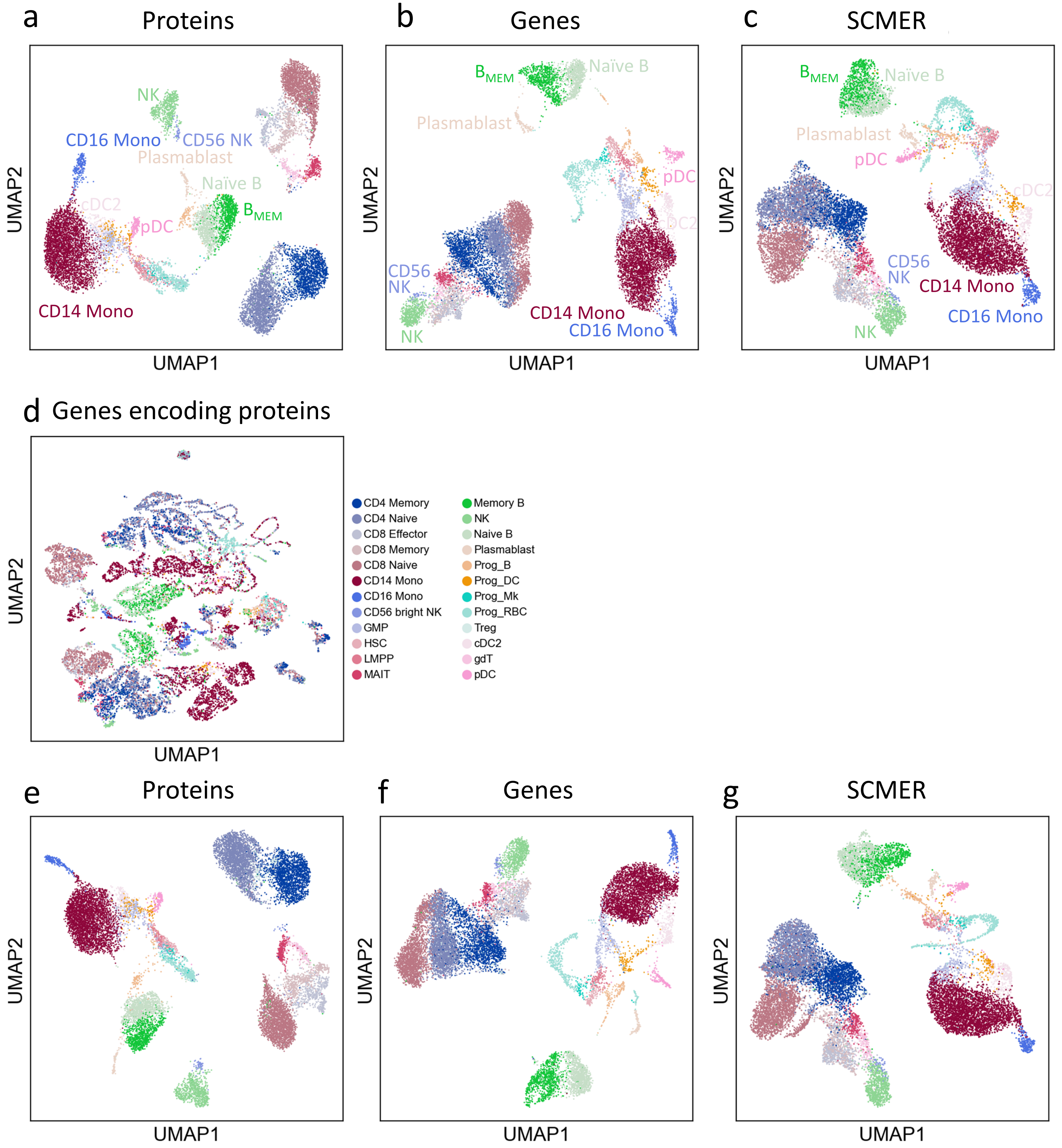


**Supplementary Figure 10:** Additional results on CITE-seq Data.
(**a-e**) UMAP embedding of original dataset using (a) protein, (b) genes, (c) SCMER selected genes, and (d) genes encoding the proteins.
(**e-g**) UMAP embedding of the validation on cell from another donor.

Supplementary Fiugre 11

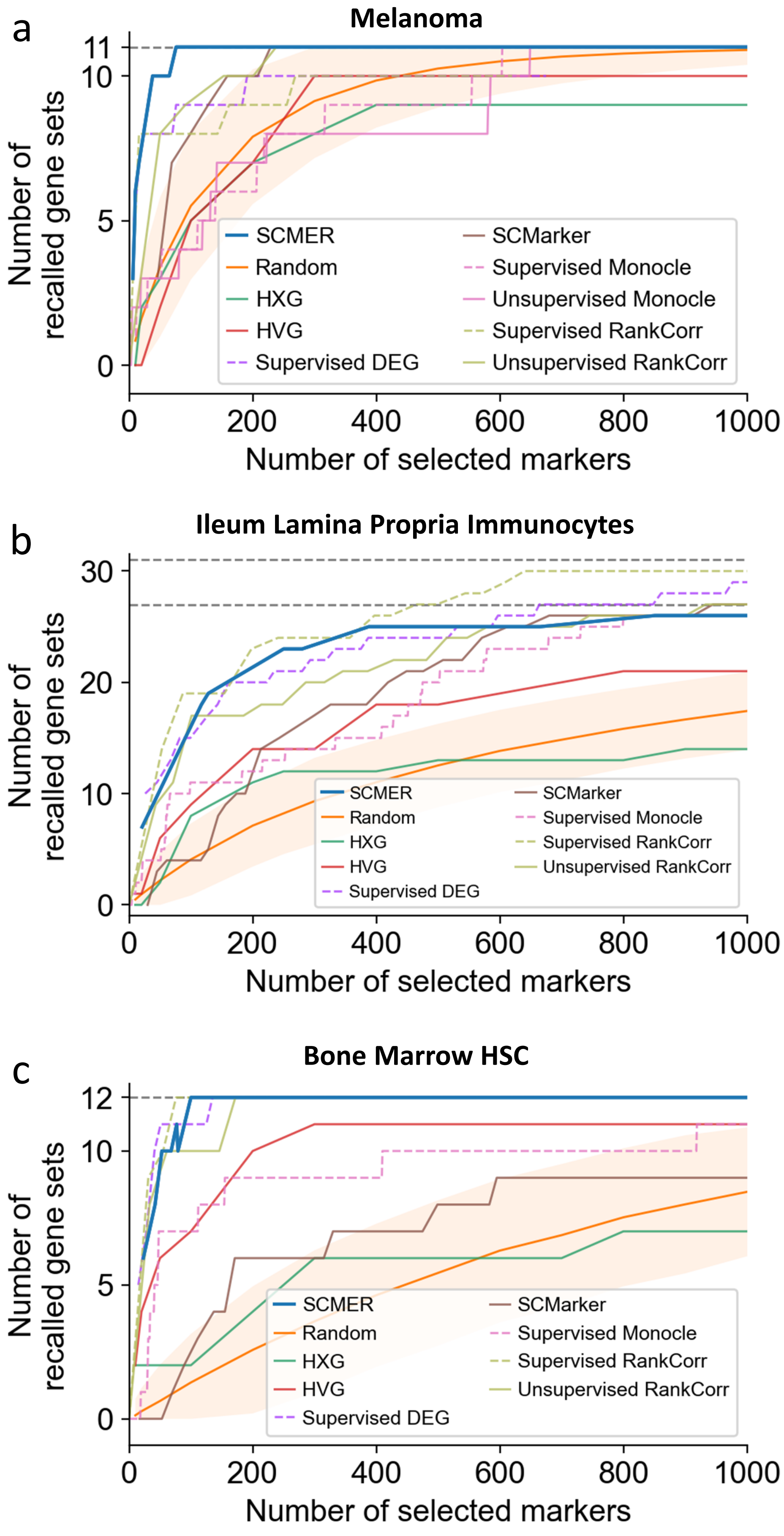


**Supplementary Figure 11:** Comparison of performances including supervised methods. (**a**-**c**) Comparison of performances of methods as in **Fig. 2c**,**3i**,**4c**, respectively, with additional supervised methods, Dashed lines represents supervised methods.

Supplementary Fiugre 12

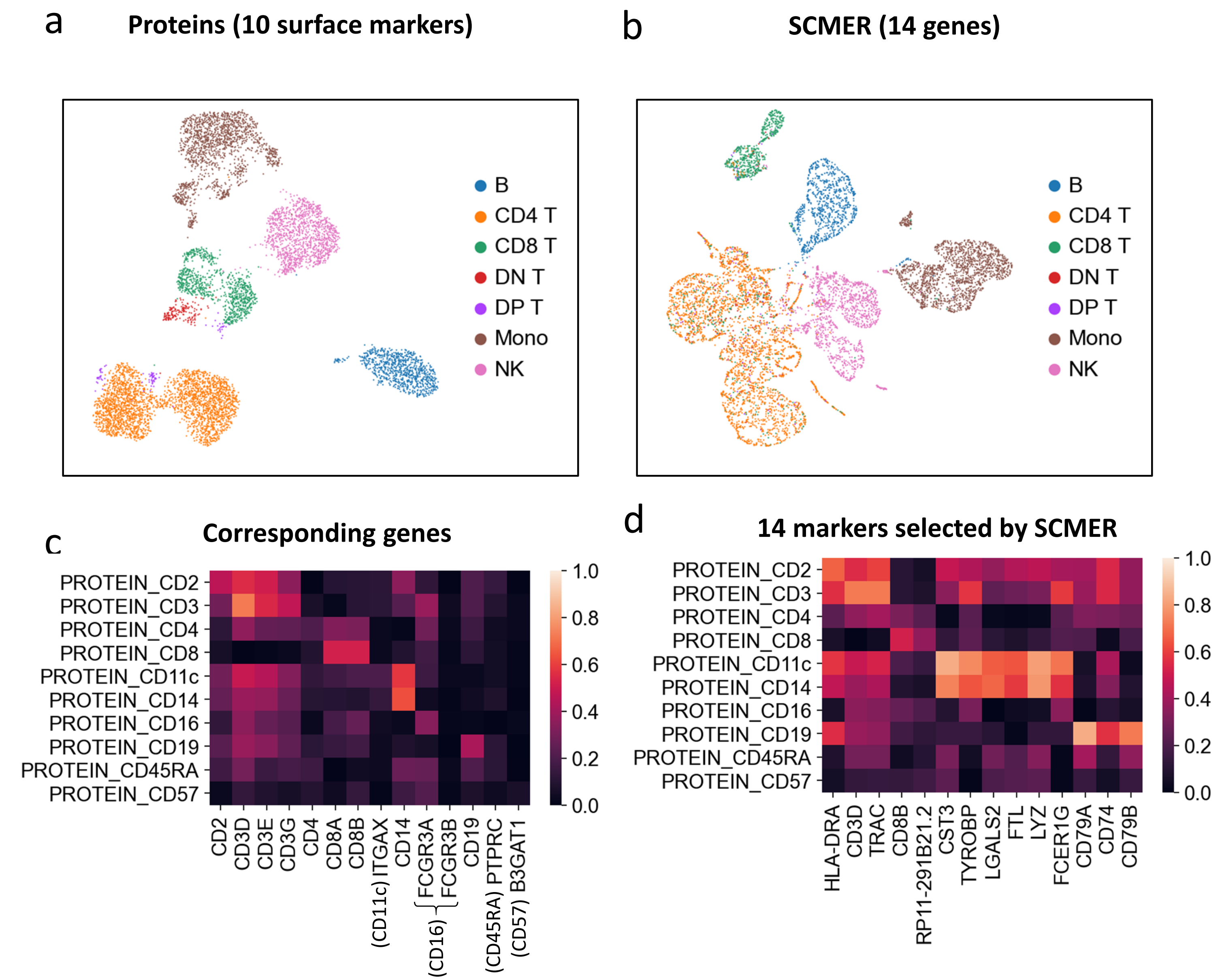


**Supplementary Figure 12:** Results on CITE-seq Data (10-protein).

(**a**) UMAP embedding of original dataset using surface markers.

(**b**) UMAP of the dataset on SCMER selected mRNA markers.

(**c**) Correlation (in absolute value) of surface markers and their corresponding genes.

(**d**) Correlation (in absolute value) of surface markers and mRNA markers found by SCMER.

### Supplementary Fiugre 13

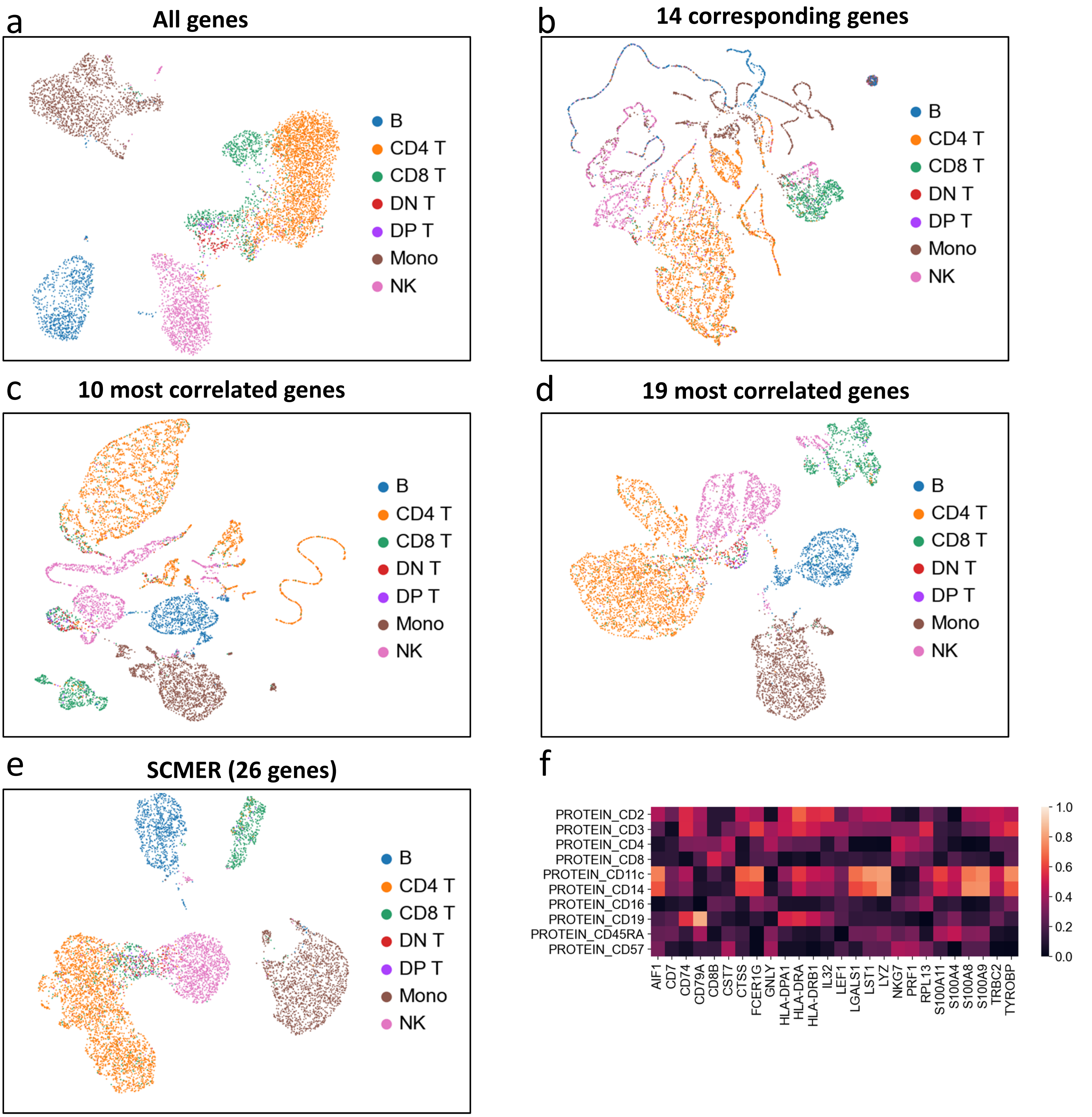


**Supplementary Figure 13:** Additional results on CITE-seq Data (10-protein).
(**a**-**e**) UMAP embedding of original dataset using (a) all genes, (b) 14 corresponding genes of the surface markers, (c) 10 most correlated genes of the surface markers, (d) 19 most correlated genes of the surface markers, and (e) 26 markers selected by SCMER.
(**f**) Correlation (in absolute value) of epitopes and 26 markers found by SCMER

### Supplementary Fiugre 14

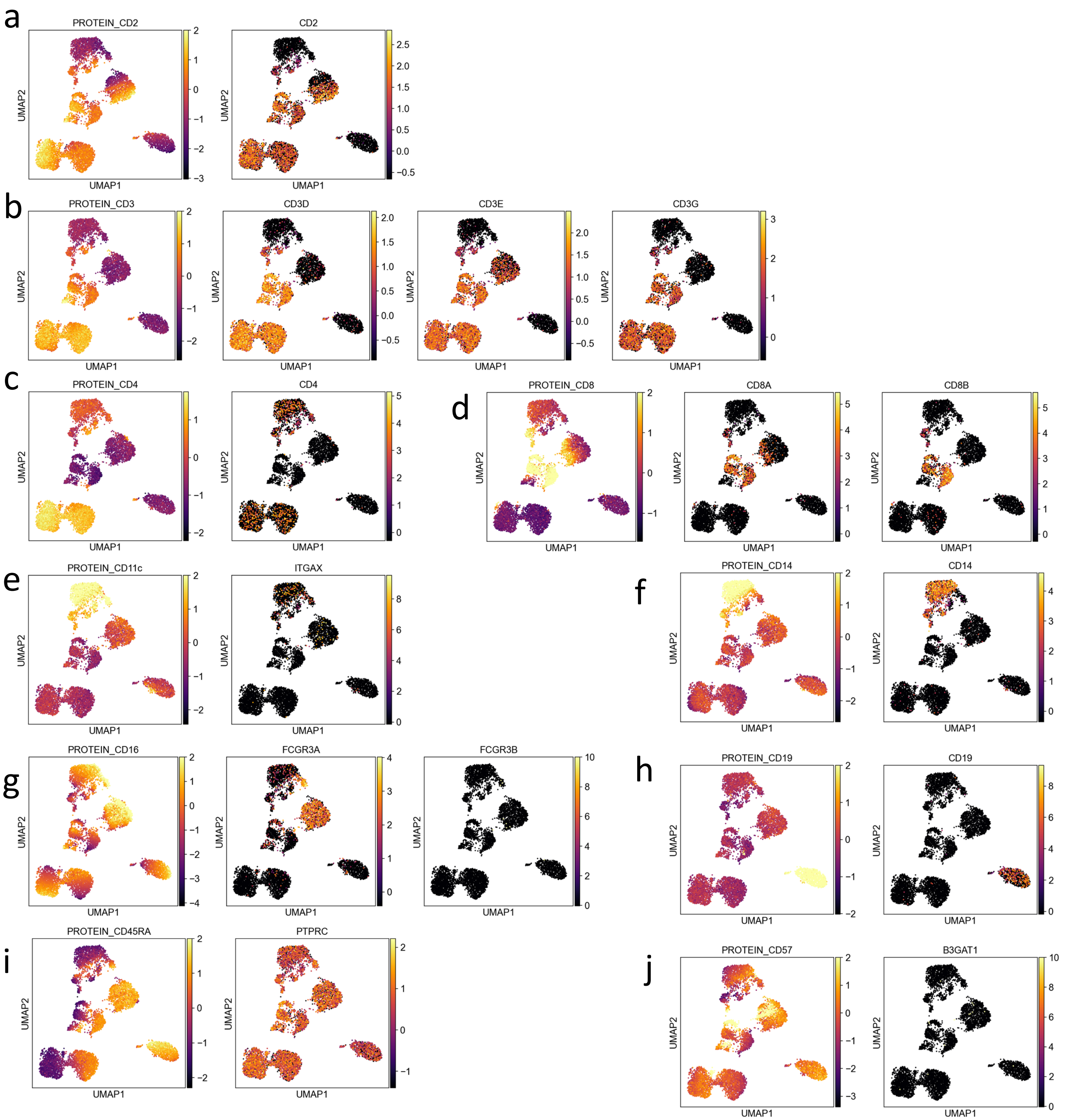


**Supplementary Figure 14:** Additional results on CITE-seq Data (10-protein). (**a**-**j**) Activity of surface markers and the genes encoding them.

Supplementary Fiugre 15

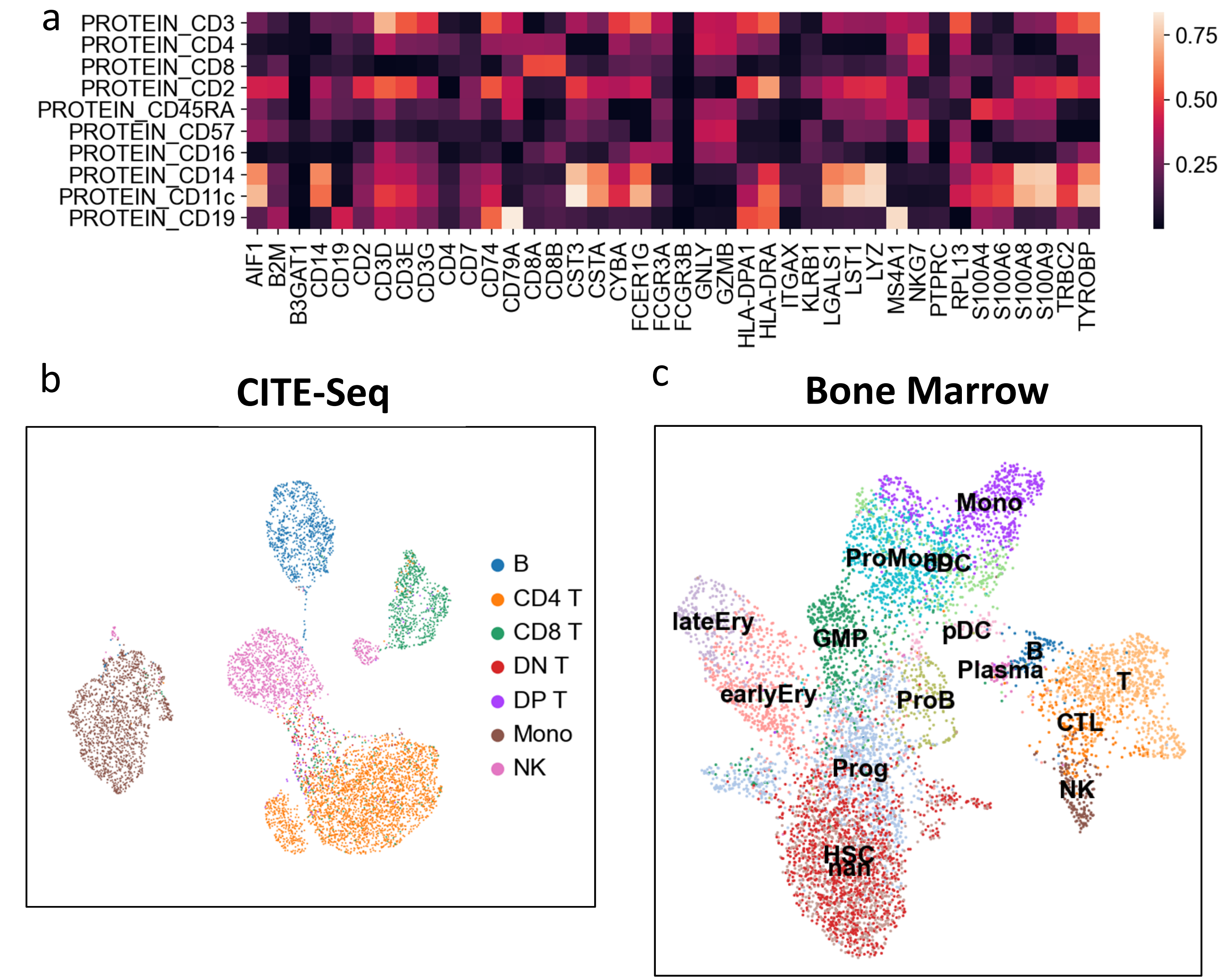


**Supplementary Figure 15:** Supervised modes of SCMER.
(**a**) Correlation of surface markers and their 14 preselected genes and 26 genes selected by SCMER accordingly on the CITE-seq data.
(**b**) UMAP of the CITE-seq data on the 40 genes in (a).
(**c**) UMAP of the Bone Marrow data on selected novel features selected by SCMER.

Supplementary Figure 16

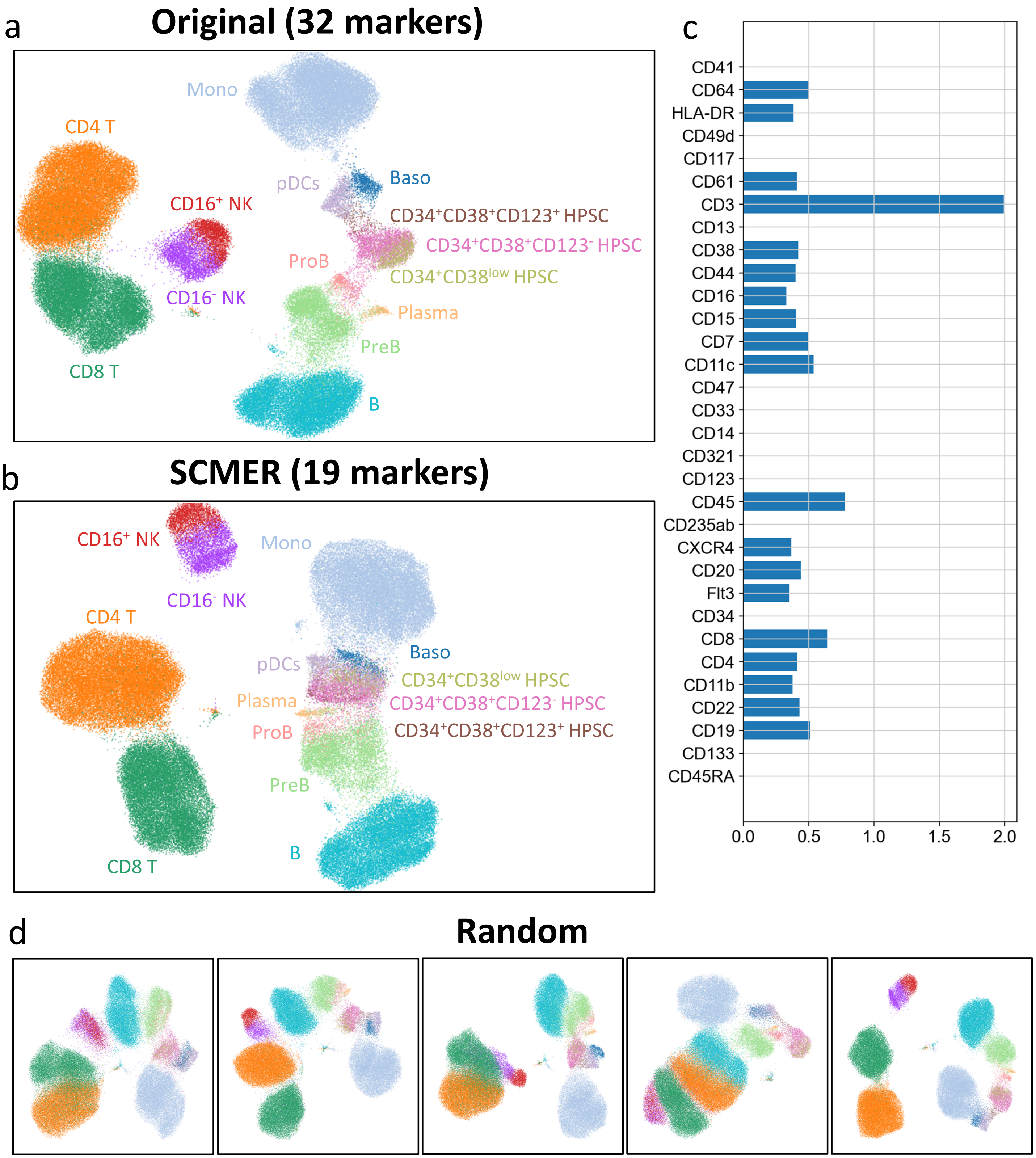


**Supplementary Figure 16:** Results on the CyTOF data.
(a) UMAP embedding of original dataset using all surface markers.
(b) UMAP of the dataset on SCMER selected mRNA markers.
(c) Weight of markers as a result of SCMER. (The weight is not used in other procedures.)
(d) UMAP of five randomly selected 19 marker panels.
